# Genetically diverse mouse models of SARS-CoV-2 infection reproduce clinical variation in type I interferon and cytokine responses in COVID-19

**DOI:** 10.1101/2021.09.17.460664

**Authors:** Shelly J. Robertson, Olivia Bedard, Kristin L. McNally, Carl Shaia, Chad Clancy, Matthew Lewis, Rebecca M. Broeckel, Abhilash I. Chiramel, Jeffrey G. Shannon, Gail L. Sturdevant, Rebecca Rosenke, Sarah L. Anzick, Elvira Forte, Christoph Preuss, Candice N. Baker, Jeffrey Harder, Catherine Brunton, Steven Munger, Daniel P. Bruno, Justin B. Lack, Jacqueline M. Leung, Amirhossein Shamsaddini, Paul Gardina, Daniel E. Sturdevant, Jian Sun, Craig Martens, Steven M. Holland, Nadia A. Rosenthal, Sonja M. Best

## Abstract

Inflammation in response to severe acute respiratory syndrome coronavirus-2 (SARS-CoV-2) infection drives severity of coronavirus disease 2019 (COVID-19) and is influenced by host genetics. To understand mechanisms of inflammation, animal models that reflect genetic diversity and clinical outcomes observed in humans are needed. We report a mouse panel comprising the genetically diverse Collaborative Cross (CC) founder strains crossed to human ACE2 transgenic mice (K18-hACE2) that confers susceptibility to SARS-CoV-2. Infection of CC x K18- hACE2 resulted in a spectrum of survival, viral replication kinetics, and immune profiles. Importantly, in contrast to the K18-hACE2 model, early type I interferon (IFN-I) and regulated proinflammatory responses were required for control of SARS-CoV-2 replication in PWK x K18-hACE2 mice that were highly resistant to disease. Thus, virus dynamics and inflammation observed in COVID-19 can be modeled in diverse mouse strains that provide a genetically tractable platform for understanding anti-coronavirus immunity.

**One Sentence Summary:** Genetically diverse mice model a spectrum of clinically relevant innate immune responses to SARS-CoV-2 infection.

## INTRODUCTION

The COVID-19 pandemic continues to pose a global threat to public health, due in part to sequential evolution of new variants of concern (VOC) selected to evade immunity generated from previous infection or vaccination. The extreme variability in patient responses to infection, ranging from asymptomatic to life-threatening illness^1–3^, remains only partially understood. In addition, persistent post-COVID health problems can be severe and include a significant risk for future morbidity and mortality^4^. The variability in disease presentation is dependent on genetic polymorphisms, age, sex and the presence of underlying conditions^1, 5, 6^, underscoring the need for research models that reflect the diverse biology and pathology of SARS-CoV-2 infection in order to provide optimal platforms for preclinical development of novel therapeutic strategies.

Type I interferon (IFN-I) is essential for control of virus replication, but its functions in COVID-19 are complex and poorly understood, with evidence for roles in both protective and pathogenic host response^7^. Success of IFN-I in controlling virus replication and orchestrating an effective inflammatory response may relate to the timing of IFN-I induction relative to peak SARS-CoV-2 replication, the intensity of IFN-I expression and the kinetics of response resolution^8^. In humans, favorable outcomes of COVID-19 have been associated with robust early IFN-I responses that are subsequently resolved along with other disease signs^9^. However, the more severe manifestations of COVID- 19, including reduced oxygen saturation, higher inflammatory responses and lower anti-viral antibody responses, have been observed in patients with either an early IFN-I response that fails to resolve^10^, or an IFN-I response that is substantially delayed relative to peak virus replication^11–14^. The roles of IFN-I in a pathogenic response may be linked to failure of control of virus replication. However, late, or sustained IFN-I responses may drive pathogenesis through failure to orchestrate an effective adaptive response, exacerbation of inflammatory monocyte responses and inhibition of tissue repair^7, 10^. In mice, loss of IFN-I signaling reduces inflammatory cell recruitment in the lung, but a direct antiviral role of IFN-I is less clear^15, 16^. Thus, additional experimental models are needed that inform how timing, magnitude, and duration of innate immunity relative to virus replication dynamics determines virus dissemination and whether inflammatory responses are protective or pathogenic.

Current mouse models of COVID-19 are important tools in understanding drivers of inflammatory responses and pathology following infection with SARS-CoV-2, but are typically generated on invariant inbred backgrounds, poorly reflecting the diversity of patient outcomes and the impact of host genetics on SARS-CoV-2 infection and response to treatment. Mice humanized for the main cellular receptor for SARS-CoV-2 entry, angiotensin converting enzyme 2 (ACE2)^17^, have been essential pre-clinical infection models^18^. The most widely used model carrying a human ACE2 gene (K18-hACE2) supports high early virus replication in lung epithelial cells, resulting in inflammatory cell infiltration, interstitial edema and focal consolidation of the lung with rapid lethality following virus dissemination to the central nervous system (CNS)^19–24^. However, this model on a standard C57BL/6J genetic background does not reflect the wide variety of COVID-19 outcomes in humans^1, 5, 6^. Indeed, in the absence of comparative models with different outcomes, it is not known what human response is being modeled by SARS-CoV-2 infection of K18-hACE2 mice specifically regarding how IFN-I response dynamics relates to control of virus replication and pathology.

To determine if incorporating genetic diversity could recapitulate the broad phenotypic variation observed in human COVID-19, we crossed the K18-hACE2 strain to the genetically distinct inbred strains A/J, 129S1/SvImJ, NOD/ShiLtJ, NZO/HILtJ, PWK/PhJ, CAST/EiJ, and WSB/EiJ that along with C57BL/6J comprise the CC founders. Together, these strains represent 90% of the genetic diversity in *M. musculus* populations^25, 26^, and therefore can be used to understand genetic regulation of complex immune responses. Notably, NOD/ShiLtJ and NZO/HILtJ mice are commonly used in studies of diabetes and metabolic dysfunction, both factors contributing to COVID-19 severity^27^. In addition to the 8 CC founder strains, BALB/cJ and DBA/2J strains were included because BALB/cJ is widely used in infectious disease modelling, and DBA/2J is a founder strain for BXD strains also used in genetic mapping^28^. Here, we show that by using these genetically diverse mice we can model both early and delayed innate immune responses associated with differing control of virus replication kinetics and pathology in the lung, and formally demonstrate an effective antiviral role for IFN-I if produced early. These mice, therefore, represent invaluable tools to understand how IFN-I and proinflammatory responses are orchestrated to control SARS-CoV-2 replication and pathology.

## RESULTS

### Diverse outcomes of SARS-CoV-2 infection in CC x K18-hACE2 F1 mice

F1 progeny of CC founder x K18-hACE2 mice (hereafter abbreviated CC x K18-hACE2) were infected via intranasal inoculation with 10^3^ plaque forming units (pfu) SARS-CoV-2 strain nCoV-WA1-2020, MN985325.1 and mice were monitored for 21 days. This virus dose is sub-uniformly lethal in the prototype K18-hACE2 strain (C57BL/6J background), resulting in 80-90% of mice reaching end-point criteria by 7 days post infection (dpi) (Fig. 1a). Infection was confirmed in all survivors by seroconversion to SARS-CoV-2 nucleoprotein. Comparative analysis of the eight CC x K18-hACE2 F1 cohorts documented a broad spectrum of weight loss and survival curves characteristic to each CC founder genotype, some of which were sex-specific. Highly sensitive strains included C57BL/6J and A/J x K18-hACE2 that lost approximately 20% of their starting weight between 4 and 7 dpi with no clear sex bias (Fig. 1a). In contrast, CC x K18-hACE2 from PWK, NZO, 129S1/J (Fig. 1b), BALB/cJ and DBA/2J (Fig. S1a, b) were comparatively resistant to clinical disease, generally losing 5-10% starting weight with ∼80% of mice surviving infection. Finally, CC x K18-hACE2 of three strains (CAST, NOD, WSB) had marked sexual dimorphism, with CAST and NOD males susceptible to lethal disease, although differences in weight loss between sexes were not apparent (Fig. 1c). Yet another phenotype was observed in WSB x K18-hACE2, where infection was uniformly lethal in females. Weight loss in males began at 6 dpi, later than other groups, and surviving males exhibited sustained weight loss in contrast to the rapid weight gain associated with recovery of other CC x K18- hACE2 cohorts (Fig. 1c).

**Figure 1:**
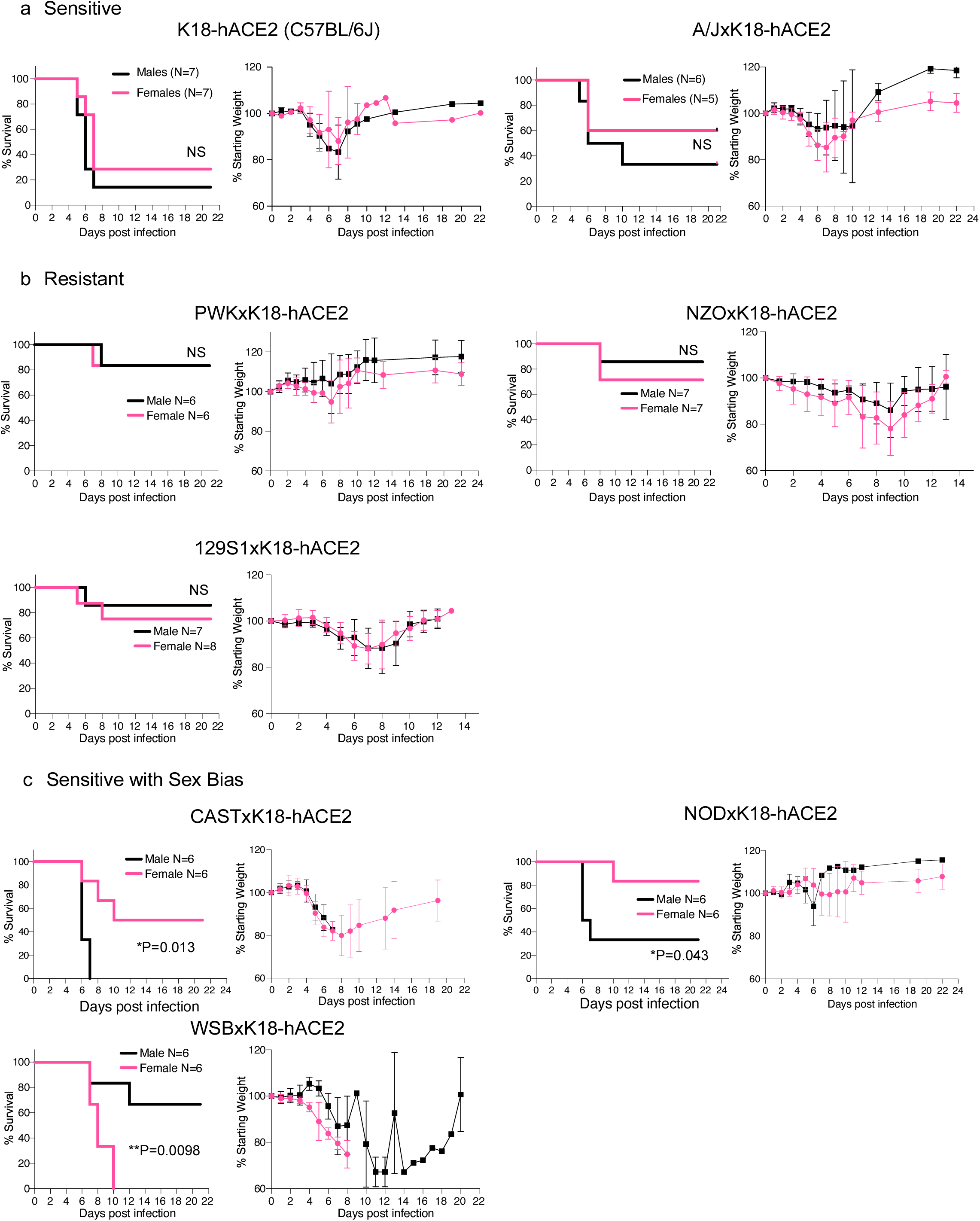
CC x K18-hACE2 mice demonstrate a range of clinical severity following infection with SARS- CoV-2. Mice were infected intranasally with 10^3^ pfu of SARS-CoV-2 and monitored for survival and weight loss. CC x K18-hACE2 were classified as: (**a)** sensitive with 50% or greater mortality and no difference between males and females, **(b)** resistant with 35% or less mortality, or **(c)** sensitive with a sex bias as having statistically significant difference between sensitivity of male and female mice. Infection was confirmed in all surviving mice by seroconversion toSARS-CoV-2 nucleoprotein (N=6-8/per sex). Graphs of percent starting weight show the mean ± SD.

### Virus replication and dissemination in CCxK18-hACE2 F1 mice

In K18-hACE2 mice, SARS-CoV-2 titers peak in the lung at 2-3 dpi and high titers of virus can be isolated from the CNS which may contribute to lethality in this model^17, 29^. Thus, virus replication kinetics were determined at 3 and 6 dpi in lung and brain of CC x K18-hACE2 cohorts (Fig. 2a, Fig. S1c). Sensitive founder strains C57BL/6J and A/J F1 progeny showed high levels of infectious SARS-CoV-2 at 3 dpi in lung homogenates that was reduced by 1-2 log10 by 6 dpi with no differences between sexes. At 3 dpi, infectious virus was generally not recovered from the CNS, but ∼50% of K18-hACE2 and A/J x K18-hACE2 had high virus burden in the CNS by 6 dpi (10^6^-10^8^ pfu/g tissue) suggesting significant virus dissemination outside of the lung. In contrast, peak lung virus titer in the most resistant PWK x K18-hACE2 was 150-fold lower than the sensitive strains at 3 dpi and was below the limit of detection at 6 dpi. Control of virus replication in PWK x K18-hACE2 was also evident in the CNS where detection of infectious virus was sporadic. Interestingly, additional CC x K18-hACE2 classed as resistant had equivalent peak viral titers in the lung to the sensitive K18-hACE2 and A/J F1 progeny, but they tended to control replication by 6 dpi to levels 1 log10 lower than sensitive mice and had less virus burden in the CNS. Finally, lung titers in (CAST, NOD and WSB) x K18-hACE2 were not different between males and females despite differences in clinical severity associated with sex. Relative expression of the K18-hACE2 transgene in lung was not different between genetic backgrounds (Fig. S2b) and ACE2 distribution was not distinguishable between sensitive (K18-hACE2) and resistant (PWK x K18-hACE2) mice by immunohistochemistry (Fig. S2c). Thus, CC x K18-hACE2 can be further stratified into a) sensitive with high sustained virus replication in lung and CNS (C57BL/6J, A/J), b) resistant with lower peak virus titer or earlier control of replication in the lung and no or low dissemination to other organs (PWK, NZO, 129S1/J), and c) sex-biased outcomes in survival independent of virus titer in the lung suggesting a sex-based difference in host response (CAST, NOD, WSB). Together, these data demonstrate that major differences in the dynamics of SARS-CoV-2 replication and host sensitivity to disease observed in humans can be modeled through use of genetically diverse mouse strains (Summarized in Table S1).

**Figure 2:**
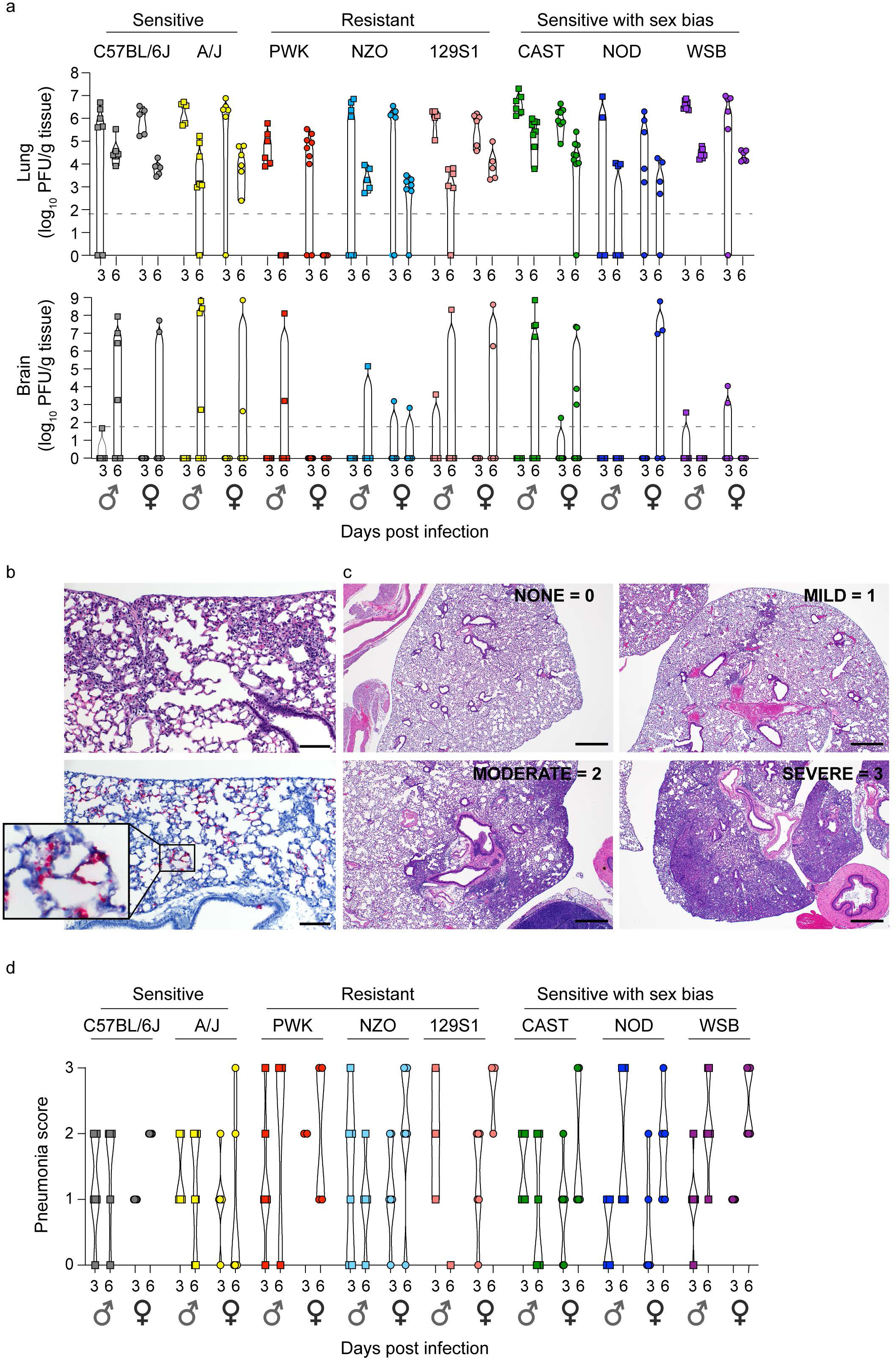
Virus replication kinetics and pathology in CC x K18-hACE2. (**a)** Virus titers (pfu/g tissue) in lung and brain of male and female mice were measured at 3 and 6 dpi. (**b)** H&E staining and *in-situ* hybridization for SARS-CoV-2 nucleic acid predominantly localizing to pneumocytes in areas of lung inflammation. Serial sections demonstrate pathology (upper) and viral RNA distribution (lower; Red). Scale bars represent 100 μm. (**c)** Examples of pathology scoring from H&E-stained lung sections. Scale bars represent 500 μm. (**d),** Lung pathology scores for CC Founder F1 males and females at 3 and 6 dpi. N = 6/time point/sex. Violin plots show all data points **(a, d)**

### Histopathology in lung and brain from SARS-CoV-2 -infected CC x K18-hACE2 F1 progeny

The general pathological changes in SARS-CoV-2-infected K18-hACE2 mice were similar to those previously described ^19, 23, 24, 30^ although varied in severity (Fig. 2b-d). Inflammatory infiltrates evident by 3 dpi included perivascular lymphocytes with alveolar septa thickened by neutrophils, macrophages, and edema. At 6 dpi, pulmonary pathology was multifocal, and consistent with interstitial pneumonia including type II pneumocyte hyperplasia, septal, alveolar and perivascular inflammation comprised of lymphocytes, macrophages and neutrophils, with alveolar fibrin and edema evident. Bronchiolar pathology was not observed in these mice. Lung pathology was classified as none, mild (rare scattered inflammatory foci), moderate (coalescing inflammatory foci) or severe (widespread, large inflammatory foci) as exemplified in Fig. 2c. Surprisingly, most resistant strains and those with sex bias tended to have higher pneumonia scores in lungs compared to the sensitive K18-hACE2 and A/J x K18-hACE2 (Fig. 2d). Lung distribution of viral RNA was limited to type I and II pneumocytes in all strains (Fig. 2b, Fig. 5b). Importantly, these data suggest that lung pathology can be exacerbated in the context of effective control of SARS-CoV-2 replication but still be associated with rapid recovery, as modeled in PWK x K18-hACE2 and NZO x K18-hACE2. In contrast, higher pathology scores associated with uncontrolled virus replication in the lung and higher sensitivity to severe disease can be modeled by in WSB x K18-hACE2.

CNS pathology was scored as present or absent, as it generally consisted of subtle inflammatory foci, including perivascular cuffing and gliosis associated with necrotic cells observed in K18-hACE2 and DBA x K18-hACE2 males (Fig. S3a-c, g). However, pathology in the CNS of CAST x K18-hACE2 was striking in that microthrombi were evident in capillaries with extensive hemorrhage in the absence of encephalitis. Although microthrombi were associated with viral RNA distribution in the same area (Fig. S3d-f), microhemorrhage was not observed as extensively in infected K18-hACE2 mice. In addition, infection with 10^3^ pfu SARS-CoV-2 was uniformly lethal in male CAST x K18-hACE2, despite these mice having some of the lowest levels of pathology in the lung at 6 dpi of all mice examined. Thus, CAST x K18-hACE2 mice may be a suitable model for examination of microthrombus formation sometimes observed in severe human COVID-19^31–33^.

### Distinct timing and amplitude of cytokine and chemokine induction in CC x K18-hACE2 F1 mice

In-depth longitudinal analyses of immune responses in patients with COVID-19 have defined immunological correlates of disease outcome^8, 10, 12, 34–44^. In addition to high early IFN responses, moderate COVID-19 is associated with increased plasma levels of a core of pro-inflammatory cytokines, chemokines and growth factors (e.g., IL-1a, IL-1ý, IFNα, IL-12p70, and IL-17A) that are effectively resolved during infection^10^. In contrast, severe COVID-19 includes sustained expression of these markers with additional signatures including IL-6, IL-10, IL-18, IL-23, TNF, and eotaxin among others suggesting that specific timing and failure to resolve inflammatory responses are important factors in disease progression^10^.

In SARS-CoV-2-infected CC x K18-hACE2, cytokines and chemokines were quantified in the serum and bronchoalveolar lavage fluid (BAL) at 3 and 6 dpi. Cytokine levels were much higher in BAL than in serum (Fig. S4a-c; individual data points are provided in Figs. S5-S7). IFNα was variably induced at 3 dpi in all crosses with K18-hACE2 (Fig. 3a and Fig. S1e). Strikingly, K18-hACE2 and CC x K18-hACE2 of A/J, CAST and NOD produced relatively low IFNα compared to PWK, NZO, 129S1/J and WSB that produced 2- to 10-fold higher levels. At 3 dpi, IFNα production was similar in males and females within each strain except for PWK x K18-hACE2 males produced significantly higher levels than females, and in DBA x K18-hACE2 where females were higher (Fig. S1d). Importantly, high early IFNα expression at 3 dpi was resolving by 6dpi in resistant ((PWK, NZO and 129S1/J) x K18-hACE2) that most effectively controlled virus replication in the lung and CNS. In comparison, low levels of IFNα were observed in K18-hACE2 mice at 3 and 6 dpi despite high early virus burden in the lung and failure to control disseminating infection by 6 dpi (Fig. 3b, c; Fig. S5). Thus, PWK x K18-hACE2 may represent a model of effective induction and resolution of IFN responses associated with viral control. A third distinct phenotype was observed in WSB x K18-hACE2 that had high IFNα expression in BAL at 3 dpi equivalent to PWK x K18-hACE2, but these mice did not control virus replication in the lung as effectively as PWK F1 mice and had sustained weight loss in surviving individuals. Indeed, WSB x K18-hACE2 produced some of the highest IFNα among all genetic backgrounds at 3 dpi (Fig. 3a) along with high levels of cytokines and chemokines (Fig. S4a-c), which may be consistent with significant weight loss observed in both sexes (Fig. 1c). In addition, WSB x K18- hACE2 had equivalent lung virus titers to K18-hACE2 mice at 3 and 6 dpi, but virus was only sporadically recovered from brain (Fig. 2a). Despite similarities in virus burden between sexes, female WSB x K18-hACE2 mice were significantly more sensitive to severe clinical disease (Fig. 1c) and lung pathology (Fig. 2d). Thus, the WSB strain background may model the situation where high IFN responses are not associated with viral control and contribute to pathogenesis.

**Figure 3:**
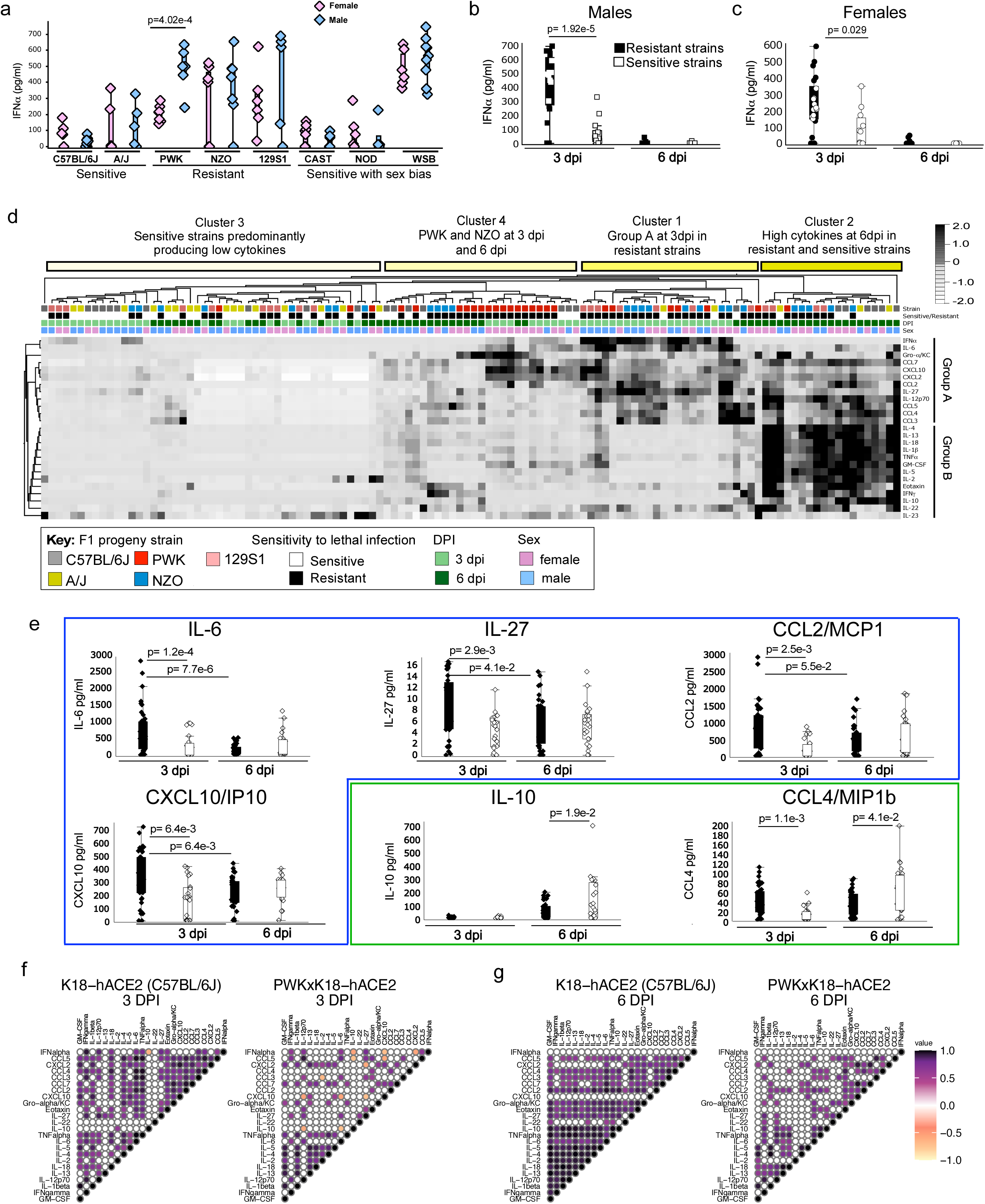
Timing of IFN-I and inflammatory cytokine responses in the lung is associated with clinical severity in CC x K18-hACE2. **(a)** Box plots of IFNα protein in BAL from CC x K18-hACE2 mice at 3 dpi. IFNα protein in BAL from **(b)** male and **(c)** female resistant (PWK, NZO and 129S1 x K18-hACE2) and sensitive (K18-hACE2 and A/J x K18-hACE2) mice at 3 and 6 dpi. **(d)** Heatmap of cytokine and chemokine responses in BAL. **(e)** Box plots of quantification of resolving group A cytokines in F1 progeny resistant to lethality (IL-6, IL-27, CCL2, CXCL10), or increasing group B cytokines in sensitive F1 progeny (IL-10, CCL4). IFNα and cytokine correlation plots for K18-hACE2 and PWK x K18-hACE2 at **(f)** 3 dpi and **(g)** 6 dpi. Box plots (drawn in Qlucore software) show individual data points with the box representing the first and third quartiles.

To further compare the relative expression of immune effectors in BAL, unsupervised hierarchical cluster analysis was performed on data from sensitive (K18-hACE2 and A/J x K18-hACE2) and three resistant strains that did not show a sex bias ((PWK, NZO, 129S1/J) x K18-hACE2). SARS-CoV-2-infected mice grouped into four general clusters each with distinct profiles of cytokine and chemokine expression. As shown in Fig. 3d, Cluster 1 was comprised of resistant mice at 3 dpi that expressed proinflammatory cytokine IL-6, TH1 cytokines (IL-12p70, IFNγ, IL-27), inflammasome associated cytokine (IL-1beta and IL-18) and chemokines (CXCL10, Gro-α/KC, CCL2, CCL3, CCL4, CCL5, CCL7 and CXCL2) (designated as Group A). At 6 dpi, some Group A cytokines were resolving (IL-6, IL-27, CCL2 and CXCL10) (Fig. 3e; Table S2; Table S3) while IL-2, IL-4, IL-5, IL-13, IL-18, IL-1b, GM-CSF, eotaxin, TNF-α, IFN-γ, and IL10 (designated as Group B) were added to the cytokine signature in resistant mice (Fig. 3d, Cluster 2). Interestingly, resistant PWK and NZO x K18-hACE2 mice generally exhibited a unique profile characterized by tempered levels of Group A and Group B along with high production of Gro-α/KC, CCL7, CXCL10, CCL2, eotaxin, and IFNγ, at 6 dpi, a timepoint coincident with the most efficient clearance of virus from the lung (Cluster 4). In contrast, sensitive strains generally failed to produce Group A cytokines at 3 dpi (Cluster 3) and if induced, production of both Group A and Group B mediators occurred at the later timepoint (Fig. 3d, Cluster 2). In contrast, sensitive F1 progeny produced significantly higher levels of IL-10 (immune suppressive cytokine) and CCL4 (monocyte and neutrophil recruiting chemokine) at 6 dpi compared to resistant F1(Fig. 3e; Table S2).

### Type I IFN signaling controls early SARS-CoV-2 replication and coordinates inflammatory responses in PWK x K18-hACE2

A correlation coefficient analysis was performed to determine the relationship between IFNα and other immune mediators, focusing on sensitive (K18-hACE2) and resistant (PWK x K18-hACE2) mice. In PWK x K18-hACE2, high levels of IFNα at 3 dpi positively correlated with a small subset of cytokines (IFNγ, IL-12p70, IL-6, IL-27 and CCL5) and negatively correlated with IL-10, CXCL10 and CXCL2 (Fig. 3f). In K18-hACE2 mice, low IFNα production correlated with low expression of these same immune mediators along with pro-inflammatory TNFα and IL-18, and immune cell recruiting chemokines (eotaxin, CCL3 and CCL4). Although IFNα expression was resolved to near basal levels in both sensitive and resistant F1 mice at 6 dpi (Fig. 3b, c) expression correlations between immune mediators increased in sensitive K18-hACE2, but generally decreased in resistant PWK x K18-hACE2 (Fig. 3g), consistent with high dysregulation in K18-hACE2.

To obtain an unbiased, global assessment of host responses in a direct comparison between resistant and sensitive mouse models, we performed Visium spatial transcriptomics on lungs from male K18-hACE2 and PWK x K18- hACE2 at 0 (naïve), 3 and 6 dpi. Confidence calling of cell types suggested that a significant proportion of Visium spots captured transcriptional signatures representing more than one cell type, predominantly endothelial cells and pneumocytes as expected in the lung (Fig. 4a, d). In K18-hACE2 mice, transcriptional changes following infection were relatively modest, with the largest change in gene signatures associated with myeloid cells, classical monocytes, and alveolar macrophages only by 6 dpi (Fig. 4b, c). In contrast, PWK x K18-hACE2 mice demonstrated dynamic changes in the lung, including a marked stromal cell response as well as recruitment of classical monocytes at 3 dpi that was resolving by 6 dpi (Fig. 4e). Evidence of T cell infiltration was also observed at 3 dpi in gene cluster 3 that contained monocytes and endothelial cells (Fig. 4d, f). In contrast, endothelial cell gene expression was reduced in representation at both 3 and 6 dpi compared to naïve lungs. However, this is likely due to increased cellular complexity of the lung at 3dpi, and a greater transcriptional signature of type II pneumocytes by 6 dpi (Fig. 4d) which may indicate cellular proliferation and repair at this time. These results demonstrate an early, orchestrated tissue-level response in PWK x K18-hACE2 mice resulting in coordinated monocyte and T cell infiltration, that was substantially delayed in the K18-hACE2 mouse model.

**Figure 4:**
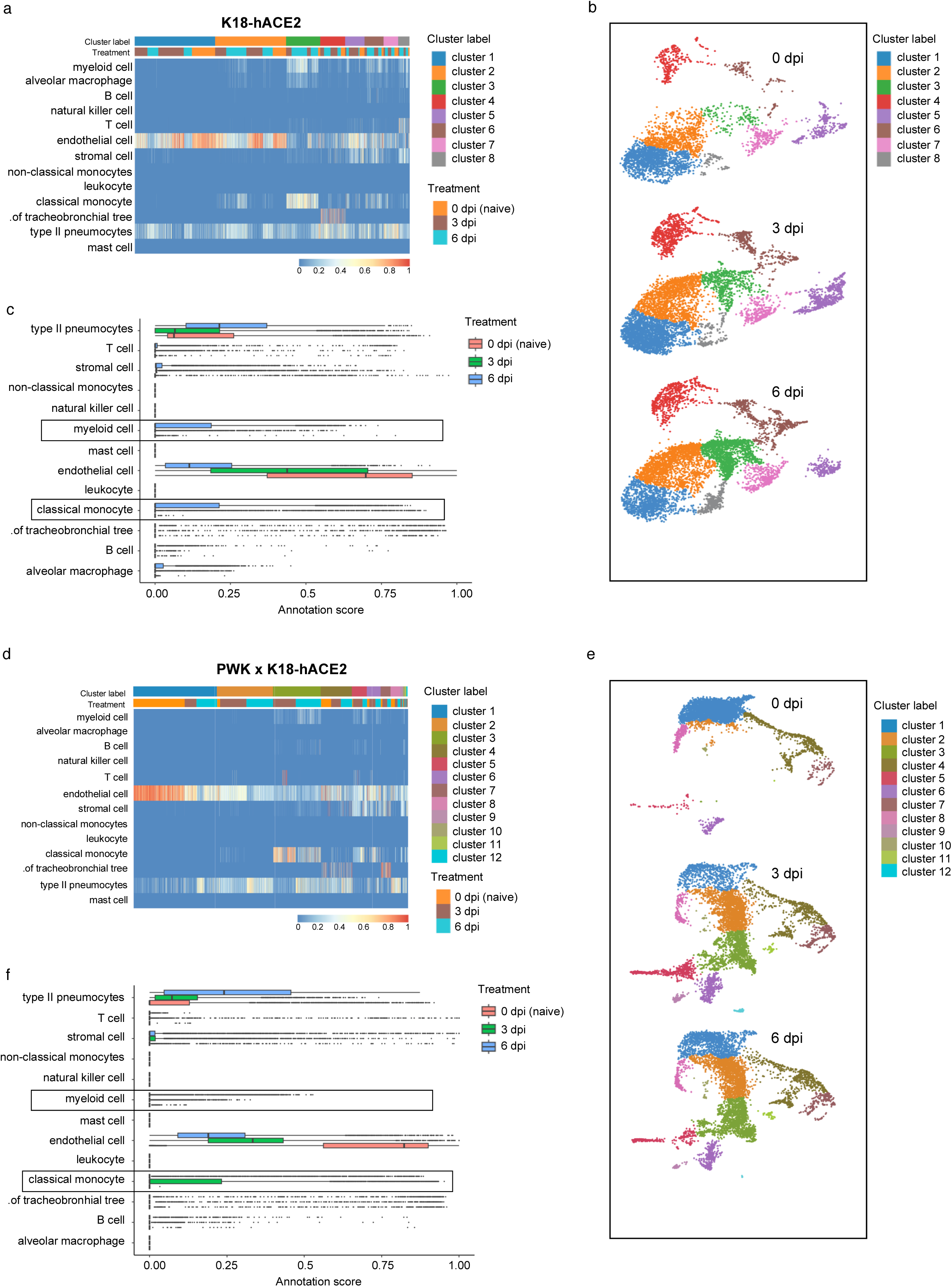
Spatial transcriptomics reveals differential innate immune signaling responses of pneumocytes, endothelial cells and monocytes in K18-hACE2 and PWK x K18-hACE2 mice. 10x Genomics Visium spatial transcriptomic analysis of lung sections from K18-hACE2 and PWK x K18-hACE2 at 0 (naïve), 3 and 6 dpi following SARS-CoV-2 inoculation. **(a, d)** Confidence scores for cell type annotations based on unsupervised cluster analysis in K18-hACE2 **(a)** and PWK x K18-hACE2 **(d)**. **(b, e)** Bar graph of cell annotations scores in K18- hACE2 **(b)** and PWK x K18-hACE2 **(e)**. **(c, f)** UMAP representation of identified transcriptional clusters in K18- hACE2 **(c)** and PWK x K18-hACE2 **(f)**.

Previous studies in C57BL/6 mice have demonstrated a role for IFNAR signaling in immune cell recruitment to the lung, but little to no role in control of SARS-CoV-2 replication or protection from clinical disease ^15, 16^. From the results obtained thus far, we suspected that low levels of IFN-I expression in K18-hACE2 at early time points enables uncontrolled virus replication but also renders this strain difficult to study the antiviral functions of IFN-I. To explore this further, expression dynamics of ISGs previously demonstrated to have direct anti-viral functions towards SARS-CoV-2^45^were examined. Like the monocyte response in PWK x K18-hACE2 mice, expression of representative ISGs *Ifi205, Zbp1, Ifit3, Ly6e, Gbp3 and Bst2* uniformly peaked at 3 dpi and was resolving by 6 dpi (Fig. 5a). ISG expression changes were generally apparent across all transcriptional clusters, suggesting that no single cell type was uniquely responsible. In contrast, ISG expression in K18-hACE2 mice was not uniformly orchestrated and was generally muted compared to PWK x K18-hACE2 mice. Low levels of *Ifi205, Zbp1 and Gbp3* expression at 3 dpi were further increasing at 6 dpi, whereas increased *Ifit3* and *Bst2* expression at 3 dpi was sustained at 6 dpi, and *Ly6e* expression was not changed over time. The dynamics of ISG expression appeared closely related to virus clearance. By RNAscope analysis for SARS-CoV-2 nucleic acid, virus was similarly distributed throughout lungs of both mouse genotypes at 3 dpi (Fig. 5b). By 6 dpi, virus distribution was unchanged in K18-hACE2 mice, but was almost completely cleared from PWK x K18-hACE2 lungs. Thus, the comparison between these highly sensitive and resistant genetic backgrounds reveals orchestrated IFN-I and monocyte responses are closely associated with control of virus replication.

**Figure 5:**
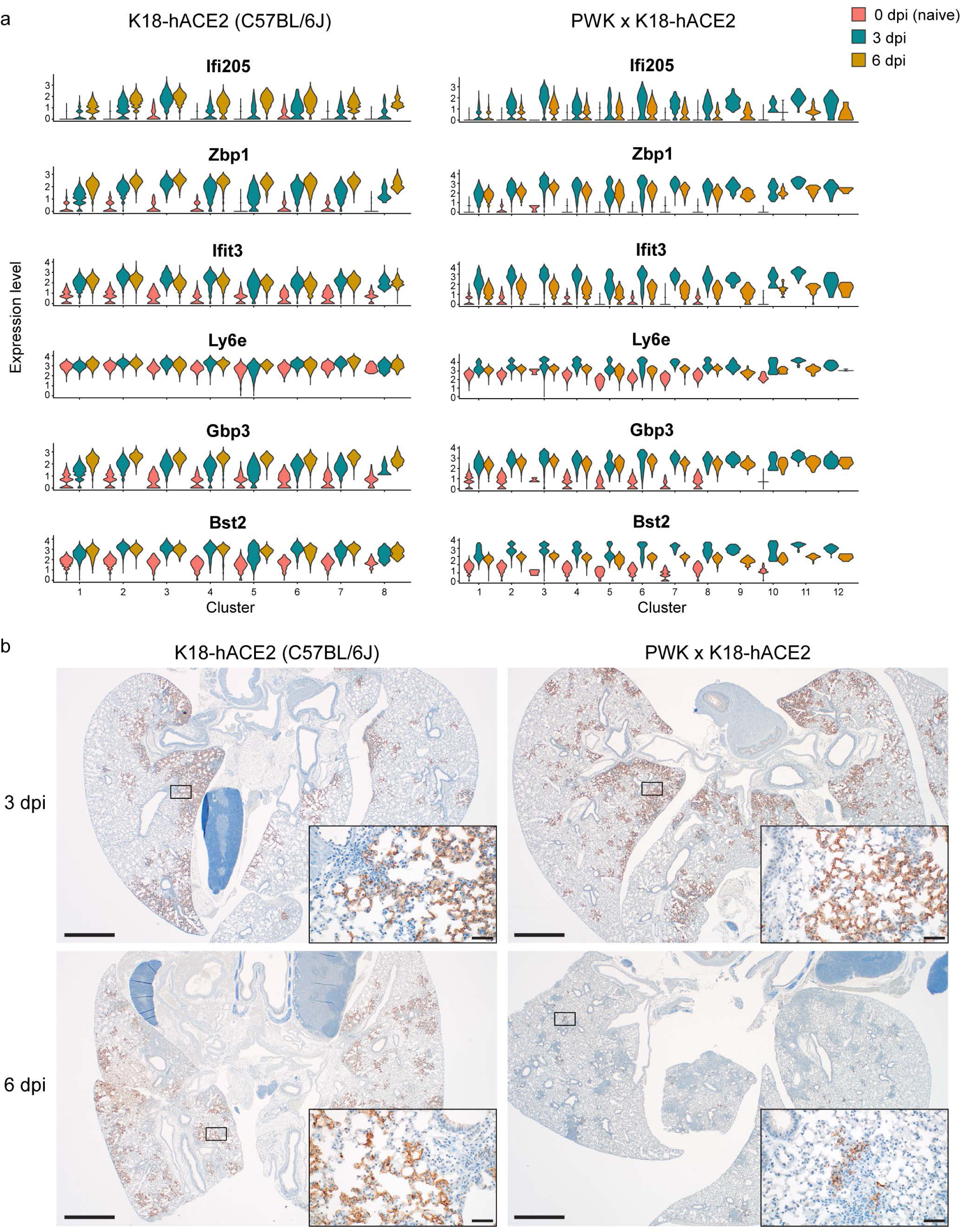
Rapid kinetics of ISG expression associated with virus clearance in PWK x K18-hACE2. **(a)** Expression levels of individual ISGs identified in Visium spatial transcriptomics analysis. **(b)** *In-situ* hybridization for SARS-CoV-2 nucleic acid in lung from K18-hACE2 and PWK x K18-hACE2 mice at 3 and 6 dpi. Scale bars represent 1000 μm; scale bars in insets represent 50 μm.

The distinct contrasts between K18-hACE2 and PWK x K18-hACE2 mice in ISG signatures, chemokine expression and viral dynamics prompted us to test the role of IFN-I by treating both genetic backgrounds with a single dose of anti-IFNAR neutralizing monoclonal antibody 24 h prior to infection. For these studies, K18-hACE2 and PWK x K18-hACE2 were inoculated with a lethal dose (10^4^ pfu of SARS-CoV-2), with male PWK x K18-hACE2 maintaining a highly resistant phenotype (Fig. S2a). Consistent with correlation coefficient analysis, neutralization of IFN-I signaling in male mice resulted in reduced expression of most inflammatory cytokines and chemokines in BAL fluid at 3 and 6 dpi regardless of genetic background (Fig. 6a). Exceptions in PWK x K18-hACE2 mice included increased IFNα and IL-12p70 at 3dpi, and no change in IFNγ and IL-18 at 6 dpi. Virus titers in lung of PWK x K18-hACE2 mice were increased by approximately 8-fold at 3dpi, with virus clearance delayed in most mice, increased neuroinvasion, and increased lethality (Fig. 6b-d). In contrast, while neutralization of IFN-I signaling resulted in a 5-fold increase in infectious titers in C57Bl/6 K18-hACE2 lung at 3dpi, it did not change kinetics of virus clearance or rates of survival (Fig. 6d). Thus, early IFN-I expression in PWK x K18-hACE2 controls peak virus loads, kinetics of virus clearance and dissemination to tissues outside of the lung. Importantly, this work definitively demonstrates the antiviral role of IFN-I to SARS-CoV-2 in vivo and illustrates the role of host genetics in determining these critical events.

**Figure 6:**
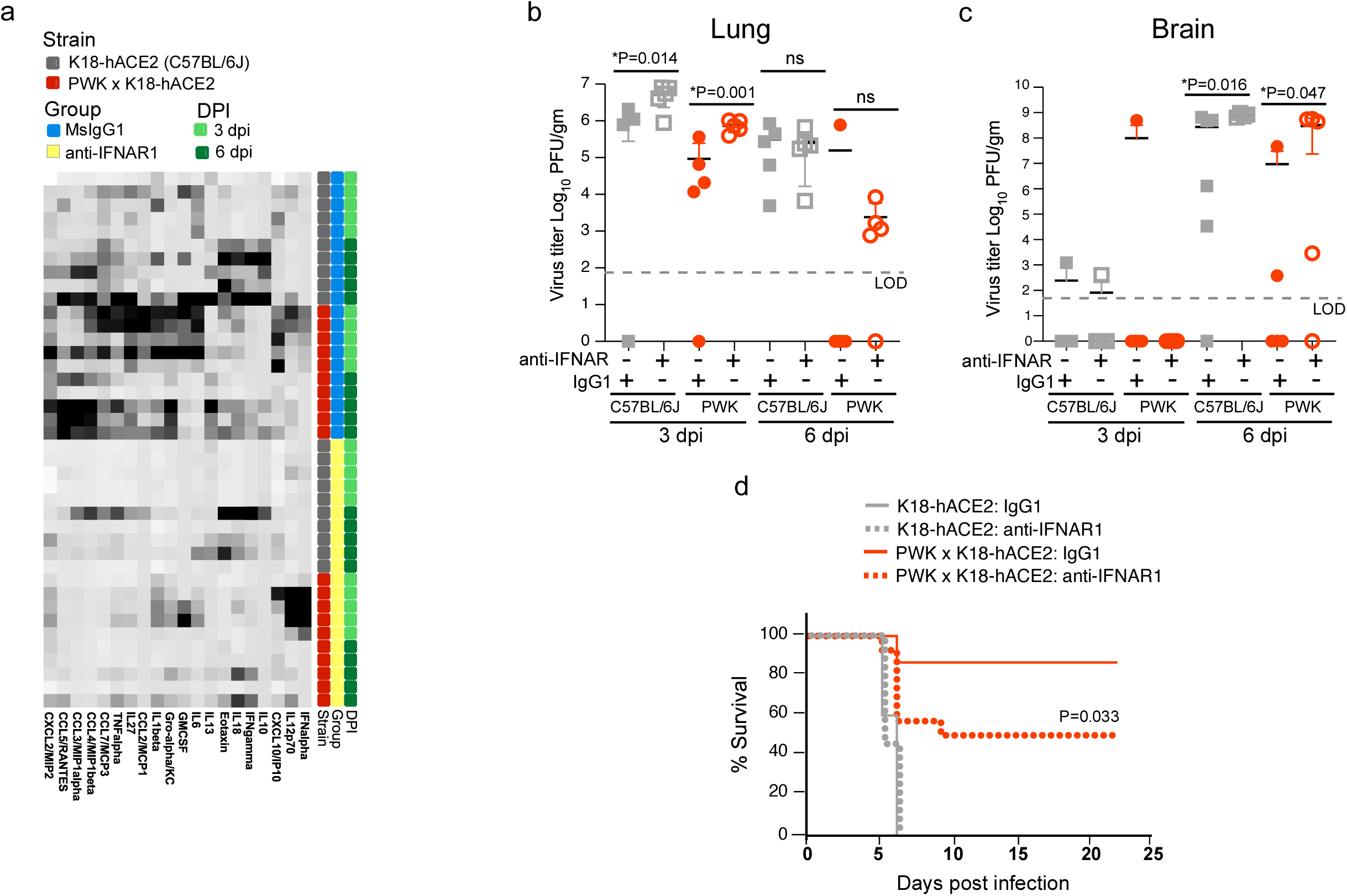
Type I IFN signaling controls early SARS-CoV-2 replication and coordinates inflammatory responses in PWK x K18-hACE2 mice. Male K18-hACE2 and PWK x K18-hACE2 mice were treated with anti- IFNAR or msIgG1 isotype control 1 day prior to intranasal inoculation with 10^4^ pfu of SARS-CoV-2. **(a)** Heatmap of cytokine and chemokine levels in BAL collected at 3 and 6 dpi. N=5/timepoint/experimental group. Virus titers (pfu/g tissue) in lung **(b)** and brain **(c)** at 3 and 6 dpi **(d).** Percent survival of mice followed daily until 21 dpi. N=10- 15/experimental group.

## DISCUSSION

Although the COVID-19 pandemic has spurred unprecedented analyses of human cohorts, the drivers of dysregulated inflammation and how they relate to virus replication kinetics in tissues are not understood. Thus, animal models that recapitulate the complexities of human COVID-19 virus replication dynamics and inflammatory responses are essential to better understand mechanisms of protection and pathogenesis. Here, we report an expanded range of SARS-CoV-2 disease phenotypes in mice using F1 progeny from the eight CC founder strains, as well as BALB/cJ and DBA/2J, crossed to K18-hACE-2 transgenic mice. The various genetic backgrounds yielded a spectrum of disease phenotypes including severe (C57BL/6J and A/J), severe with sex-bias (CAST/EiJ, NOD/ShiLtJ and WSB/EiJ), and resistant mice exhibiting relatively high pathology but non-lethal disease (PWK/PhJ, NZO/HILtJ, 129S1/SvImJ, DBA/2J and BALB/cJ). Variability in outcomes of SARS-CoV-2 infection in these mouse models was associated with distinct dynamics of innate immunity. Herein, control of virus replication in the lung and host resistance to disease can be independent from the early peak titer, but it was closely associated with a phased amplification and resolution of proinflammatory mediators associated with rapid control of virus replication in lung and prevention of virus dissemination to other organs. These favorable outcomes of COVID-19 are most clearly modeled by examining SARS-CoV-2 infection of PWK and NZO x K18-hACE2. In contrast, infection of C57Bl/6 or A/J x K18-hACE2 represent a model of inefficient IFN expression in the lung, failed host control of virus replication and dysregulated proinflammatory responses^8, 10, 43, 46–48^. Collectively, the CC x K18-hACE-2 panel represents a genetically diverse population and reflect a broader range of host-viral interactions that influence disease outcomes. Importantly, this panel provides much needed tools to further determine the roles of specific cell types and signaling pathways in orchestrating protective versus pathogenic immunity.

Early events surrounding the production of antiviral IFNs are key determinants in human outcomes of COVID-19, but a mechanistic understanding is lacking. Evidence linking IFN-I responses to dysregulated immunity and disease severity includes genetic lesions in pattern recognition receptor signaling molecules^3^, production of anti-IFN autoantibodies^12^, and patterns of IFN-I and ISG expression in respiratory tissues^49^. However, these attributes are linked to subsets of patients with severe COVID-19, which leaves open questions regarding how antiviral functions of IFNs are balanced with inflammation and why some people fail to mount effective antiviral responses despite high viral load or high IFN-I expression. The CC x K18-hACE2 mice revealed that IFN-I responses in lungs of K18- hACE2(C57Bl/6J) are relatively low compared to other specific genetic backgrounds, which may explain why ablation of IFN-I signaling was associated with relatively small effects in this model (this study, and^15, 16^). In contrast, the PWK x K18-hACE2 infection model represents the outcome where IFN-I is directly linked to control of peak virus loads, magnitude of inflammatory cytokine expression independent of virus burden, regulation of adaptive immunity, virus clearance, and control of virus dissemination to tissues outside of the lung. It is curious then that IFN-I expression was highest in BAL fluid of WSB x K18-hACE2 that had high early virus burden in the lung with delayed clearance similar to mice with low IFN-I expression (C57Bl/6J-K18-hACE2). Thus, by comparison, viral dynamics, and pathology in WSB x K18-hACE2 mice provide a model of high IFN-I driving pathology combined with inefficient antiviral functions, as these mice had some of the highest lung pathology scores. Interestingly, a survey of immune cell populations in CC founder mice demonstrated that unchallenged WSB mice have the highest percentages of plasmacytoid dendritic cells (pDCs)^50, 51^, critical producers of IFN-I in blood and tissues. It is possible that the cellular origin of IFN-I in lungs of SARS-CoV-2-infected PWK and WSB x K18- hACE2 mice is different, that pDC function in the WSB background is dysregulated as is observed in severe COVID-19 ^52^, or that high IFN-I expression and resulting pathology or inflammatory cell milieu inhibits adaptive immunity in WSB mice. Further insight will be gained by comprehensively comparing immunological and transcriptional responses of PWK and WSB x K18-hACE2 mice following challenge with SARS-CoV-2. Importantly, virus dissemination to the CNS does not consistently occur in PWK or WSB genetic backgrounds suggesting that these two new models are particularly valuable in understanding immune cell dynamics in the lung and should be explored for their potential to model post-acute events including sequalae of long-COVID.

The remarkedly different patterns of ISG expression in lungs of K18-hACE2 and PWK x K18-hACE2 mice begins to clarify how IFN-I signaling can fail to be well orchestrated and effective. We examined canonical ISGs with demonstrated antiviral functions towards SARS-CoV-2 (e.g., Bst2/tetherin and ZBP1^45^, as well as Gbp proteins that SARS-CoV-2 VOC have specifically evolved resistance to^53^. Prompt and widespread gene expression was tightly associated with efficient control of virus replication in PWK x K18-hACE2 lungs. However, while ISG expression was increased at 3 dpi of K18-hACE2 mice, it failed to effectively control virus replication resulting in further increases in host gene expression. We speculate that this sustained ISG expression in the presence of relatively high virus burden may provide the environment that facilitates virus evolution to escape individual ISGs. As expected, IFN-I was not solely required for control of virus replication in PWK x K18-hACE2 mice as IFNAR blocking experiments did not result in a rebound of virus titers to the same levels as K18-hACE2 or 100% lethality. However, the PWK genetic background will enable experimental models to empirically determine how antiviral mechanisms are coordinated to control replication, including type III IFNs that have been shown to correlate with virus control in humans^49^.

One limitation of the study is the degree of variability observed in the data likely due to the sublethal virus dose used. However, as exemplified in PWK x K18-hACE2, this was an important element of the experimental design because it enabled observation of host resistance by not overwhelming initial host defenses. Another limitation is the use of heterozygous F1 offspring because important phenotypes could be due to either or both parental alleles, making it difficult to map the genetic loci responsible for a given phenotype. However, K18-hACE2 mice were used as a common parent for all F1s and therefore, any differences observed in disease outcome is due to genes received from the unique parental strain (CC founders, BALB/cJ or DBA/2).

In summary, the phenotypes in diverse mice support a role for the innate immune system in determining COVID-19 severity^3, 8, 46–48^ and can be used to address key knowledge gaps, including mechanisms of innate immune control of virus replication, defining events needed for a well-orchestrated inflammatory response independent of early viral burden, molecular mechanisms of sex-dependent disease severity, and longer-term implications for tissue repair^54^ and lung function. Taken together, these observations demonstrate that use of host genetic variation in mice can model different outcomes of SARS-CoV-2 infection, addressing a major deficiency in the toolkit required to combat COVID-19.

## MATERIALS AND METHODS

### Ethics statement

Animal study protocols were reviewed and approved by the Institutional Animal Care and Use Committee (IACUC) at Rocky Mountain Laboratories (RML), NIAID, NIH in accordance with the recommendations in the Guide for the Care and Use of Laboratory Animals of the NIH. All animal experiments were performed in an animal biosafety level 3 (ABSL3) research facility at RML. Standard operating procedures for work with infectious SARS-CoV-2 and protocols for virus inactivation were approved by the Institutional Biosafety Committee (IBC) and performed under BSL3 conditions.

### Virus preparation

SARS-CoV-2 (USA_WA1/2020) from University of Texas Medical Branch (Vero passage 4) was propagated on Vero cells cultured in DMEM supplemented with 10% fetal bovine serum, 1 mM L-glutamine, 50U/ml penicillin and 50ug/ml streptomycin. Culture supernatants were collected at 72 hpi, aliquoted and stored at -80°C.

### Mice, virus infection, and in vivo block of IFNAR1

CC founder x C57BL/6J -K18-hACE2 F1 were provided by The Jackson Laboratories and include the following: B6.Cg-Tg (K18-ACE2)2PrImn/J), 034860; (A/J x B6.Cg-Tg(K18-ACE2)2Prlmn/J)F1/J, 035940; (PWK/PhJ x B6.Cg-Tg(K18-ACE2)2Prlmn/J)F1/J, 035938; (NZO/HlLtJ x B6.Cg-Tg(K18-ACE2)2Prlmn/J)F1/J, 035936; (129S1/SvImJ x B6.Cg-Tg(K18-ACE2)2Prlmn/J)F1/J, 035934; (CAST/EiJ x B6.Cg-Tg(K18-ACE2)2Prlmn/J)F1/J, 035937; (NOD/ShiLtJ x B6.Cg-Tg(K18-ACE2)2Prlmn/J)F1/J, 035935; (WSB/EiJ x B6.Cg-Tg(K18- ACE2)2Prlmn/J)F1/J, 035939; (BALB/cJ x B6.Cg-Tg(K18-ACE2)2Prlmn/J)F1/J, 035941; (DBA/2J x B6.Cg-Tg(K18-ACE2)2Prlmn/J)F1/J 035943. Six- to 12-week-old male and female mice were inoculated by the intranasal route with 10^3^ pfu of SARS-CoV-2 in a volume of 50ul PBS (Gibco). Prior to inoculation mice were anesthetized by inhalation of isoflurane. Mice were monitored daily for clinical signs of disease and weight loss. In IFNAR1 blocking experiments, male K18-hACE2 and PWK x K18-hACE2 mice were inoculated intraperitoneally anti- IFNAR1 (MAR1-5A3, BioCell) or msIgG1 isotype control (MOPC-21, BioCell) 1 day prior to intranasal inoculation with 10^4^ pfu of SARS-CoV-2.

### Virus titration

Infectious virus in lung and brain tissue was quantified by plaque assay. Tissues were collected in 0.5ml of DMEM containing 2% FBS, 50U/ml penicillin and 50ug/ml streptomycin and immediately frozen. Tissue samples were weighed, and then homogenized using 5mm steel beads and TissueLyzer II high-speed shaker (Qiagen). Ten-fold serial dilutions of homogenates were prepared in duplicate and used to inoculate Vero cells grown in 48-well tissue culture plates. Following 1 hour incubation at 37°C, the cells were overlayed with 1.5% carboxymethyl cellulose (CMC) in MEM and incubated at 37°C for 3-4 days. Cells were then fixed in 10%formalin and plaques were visualized by staining with 1% crystal violet diluted in 10% ethanol.

### Measurement of SARS-CoV-2-specific IgG

Sera were collected at 21 dpi from mice that survived SARS-CoV-2 infection. SARS-CoV-2 spike protein-specific IgG was measured using SARS-CoV2 spike protein serological IgG ELISA kit (Cell Signaling Technology) per manufacturer’s instructions.

### Multiplex cytokine/chemokine analysis

Sera were collected from SARS-CoV-2-infected mice at 3 and 6 dpi by centrifugation of whole blood in GelZ serum separation tubes (Sarstedt). BAL samples were recovered by insufflation of lungs with 1 ml sterile PBS followed by aspiration to collect ∼0.5ml volume of fluid. SARS-CoV-2 in sera and BAL fluid was inactivated by using γ- irradiation (2 MRad) and removed from the BSL3 laboratory. Cytokine concentrations in serum and BAL were measured using a 26-Procartaplex mouse cytokine/chemokine panel (ThermoFisher EPX260-26088-901) combined with a Simplex IFN*a* assay (ThermoFisher EPX01A-26027-901). Samples were run on a Luminex Bio-Plex 200 system with BioPlex Manager 6.1.1 software. Data represented are from samples collected from individual mice.

### Histopathology and *in situ* hybridization

Tissues were fixed in 10% neutral buffered formalin for a minimum of 7 days with 2 changes according to IBC-approved standard operating procedure. Tissues were processed with a Sakura VIP-6 Tissue Tek, on a 12-hour automated schedule, using a graded series of ethanol, xylene, and PureAffin. Embedded tissues were sectioned at 5 μm and dried overnight at 42°C prior to staining with hematoxylin and eosin. Chromogenic detection of SARS-CoV-2 viral RNA was performed using the RNAscope VS Universal AP assay (Advanced Cell Diagnostics Inc.) on the Ventana Discovery ULTRA stainer using a SARS-CoV-2 specific probe (Advanced Cell Diagnostics Inc. cat. 848569). *In situ* hybridization was performed according to manufacturer’s instructions. ACE2 immunoreactivity was detected using R&D Systems AF933 at a 1:100 dilution. The secondary antibody is the Vector Laboratories catalog number MP-7405 ImmPress anti-goat polymer. The tissues were then processed for immunohistochemistry using the Discovery Ultra automated stainer (Ventana Medical Systems) with a ChromoMap DAB kit (Roche Tissue Diagnostics cat. 760-159).

### Visium spatial transcriptomics and processing

Formalin-fixed, paraffin-embedded samples were sectioned directly onto Visium Spatial for FFPE Gene Expression Mouse Transcriptome slides (6.5 X 6.5 mm) and processed for sequencing, according to the manufacturer’s procedure (10X Genomics, Pleasanton, CA). Resulting libraries were sequenced as 28/10/10/50 base reads on the NovaSeq instrument using the S2 Reagent kit in paired-end mode (Illumina, San Diego, CA). Fastq files were processed with Spaceranger-1.4.2 pipeline. Processed data were then analyzed on R studio (2021.09.1, Build 372) with Seurat (v4.0)^55^. We trimmed spots with zero count and normalized using SCtransform function^56^. Then data were integrated using IntegrateAnchor method^57^. A total of 2000 highly variable genes identified by the SCtransform function in the Seurat^55^package were used for principal component analysis (PCA)-based dimensionality reduction with RunPCA. Uniform Manifold Approximation and Projection (UMAP) with a resolution of 0.05 was utilized to visualize single-cell clustering using principal components (PCs) 1 to 30.

### Quantitative real-time PCR

Total lung RNA was extracted from homogenized lysates using the *mir*Vana miRNA Isolation Kit (Invitrogen cat. AM1560) following the manufacturer’s protocol for total RNA isolation. Total RNA samples were quantified using a Nanodrop spectrophotometer (Thermo cat. ND-2000C), and reverse transcribed to cDNA using the SuperScript IV VILO Master Mix with ezDNase enzyme (Invitrogen cat. 11766050) following the manufacturer’s instructions. Expression of mouse *Ace2*, mouse *Gapdh*, and human ACE2 were quantified by qPCR using the ViiA7 Real-Time PCR System (Thermo cat. 44535545). TaqMan probes targeting mouse *Gapdh* (Thermo probe ID Mm99999915_g1) and human ACE2 (Thermo probe ID Hs01085333_m1) were used in conjunction with the TaqMan Fast Advanced Master Mix (Thermo cat. 4444556) and the ViiA7 Real-Time PCR System with QuantStudio Software (Thermo cat. 44535545). Samples were run in technical triplicate in 10ul reaction volumes on a 384-well plate, and hACE2 transcript abundance values were normalized to m*Gapdh* in each sample and relative expression was calculated using a reference lung sample (K18-hACE2) and the delta delta-Ct method.

### Statistical analysis

Comparison of survival curves in males versus females for each strain was performed using the log-rank (Mantel-Cox) test. One-way analysis of variance (ANOVA) with Tukey’s multiple comparison posttest was used to compare cytokine and chemokine levels of BAL and serum. Differences between groups were considered significant at a *P* value of <0.05. All statistical analyses were performed with graphPad Prism 8.0 (GraphPad Software) or Qlucore Omics Explorer Version 3.7 (Qlucore AB). Correlation analyses were performed by computing Spearman’s coefficients and visualized using ggcorrplot. Correlation tests with *P* < 0.001 were displayed.

### Data Availability

All data are available without restrictions. Visium RNA-seq data that support the findings of this study have been deposited in Gene Expression Omnibus (GEO) public database with the accession code XXX.

## Supplementary Materials

**Supplemental Figure 1:**
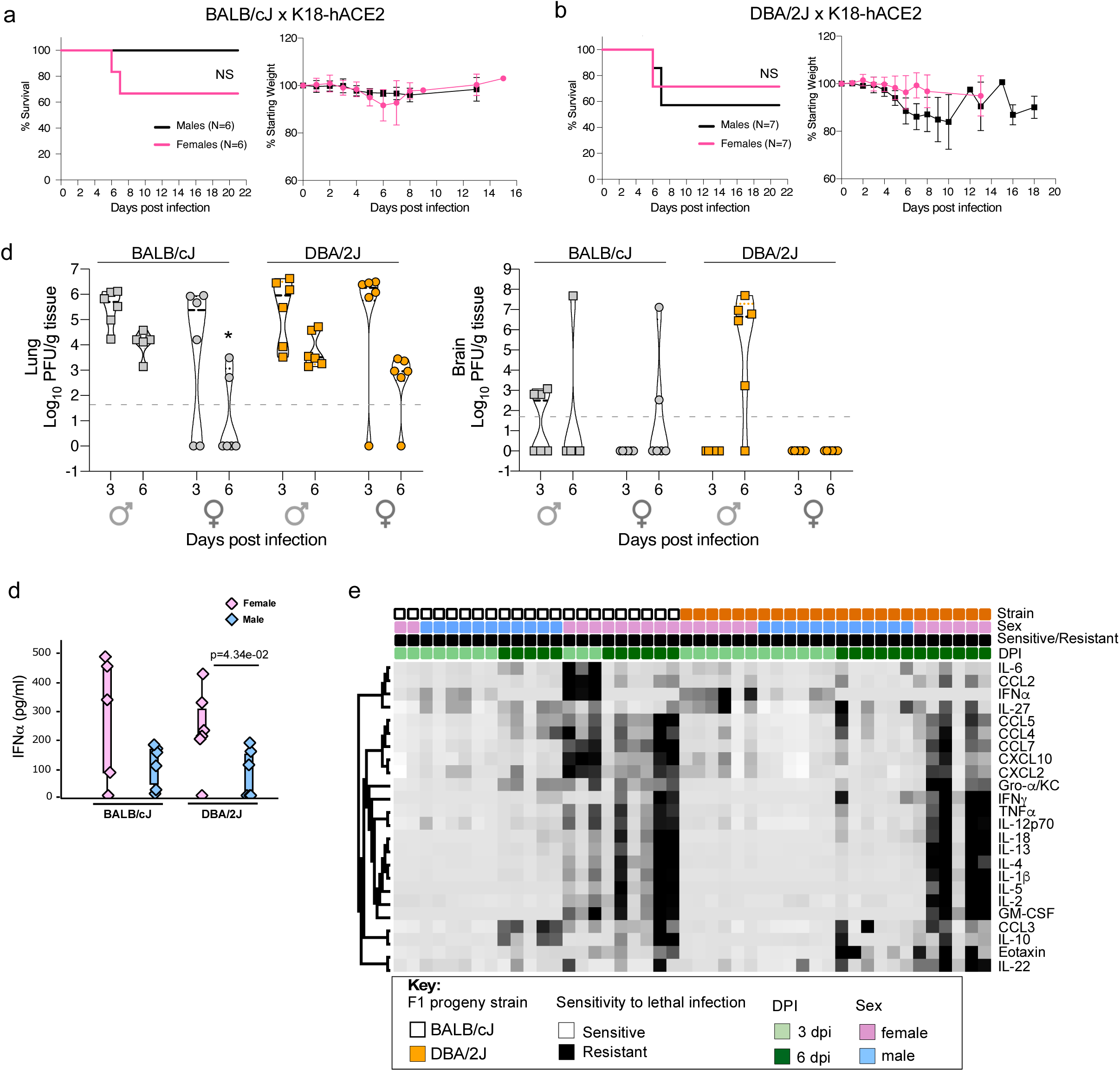
Survival, weight loss and SARS-CoV-2 replication kinetics in BALB/cJ x K18-hACE2 or DBA/2J x K18-hACE2 F1 progeny. (**a)** Survival and weight loss in BALB/cJ x K18-hACE2. (**b)** Survival and weight loss in DBA/2Jx K18-hACE2. (**c)** Virus titers (pfu/g tissue) in lung and brain of male and female mice were measured at 3 and 6 dpi. (**d**) IFNα protein in BAL from BALB/cJ and DBA/2J x K18-hACE2 mice at 3 dpi. **(e)** Heatmap of cytokine expression in BAL.

**Supplemental Figure 2:**
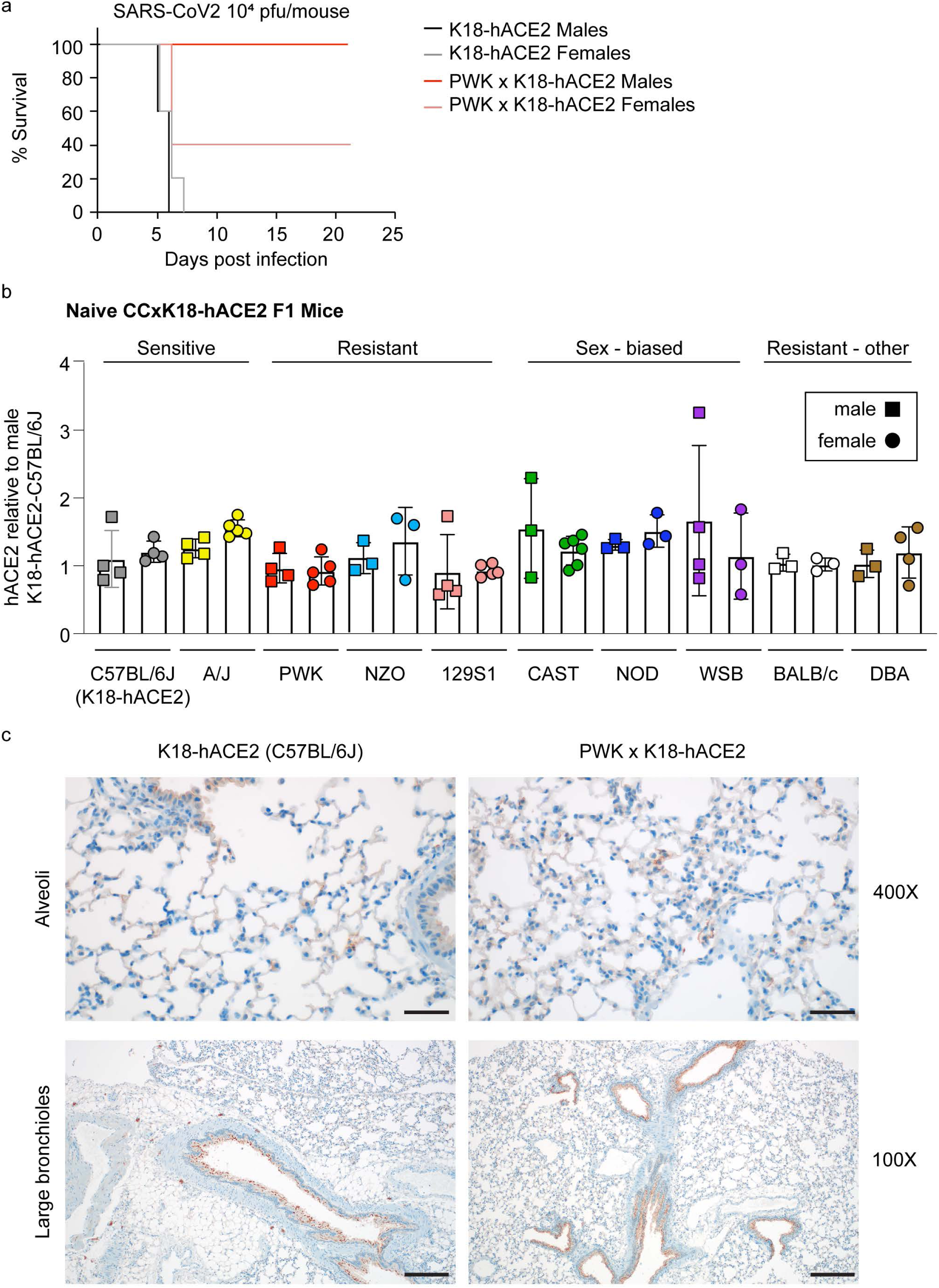
Expression of ACE2 in lungs of CC x K18-hACE2 F1 progeny. **(a)** Survival curve of K18-hACE2 and PWK x K18-hACE2 male and female mice inoculated intranasally with 10^4^ pfu of SARS-CoV-2 and monitored for survival. **(b)** Gene expression specific for *hACE2* was measured by real-time RT-PCR and expressed relative to that in male and female K18-hACE2 (C57BL/6J) mice. (**c)** IHC staining for ACE2 in K18- hACE2 and PWK x K18-hACE2 mice. Scale bars represent 50 μm (400X); 200 μm (100X).

**Supplemental Figure 3:**
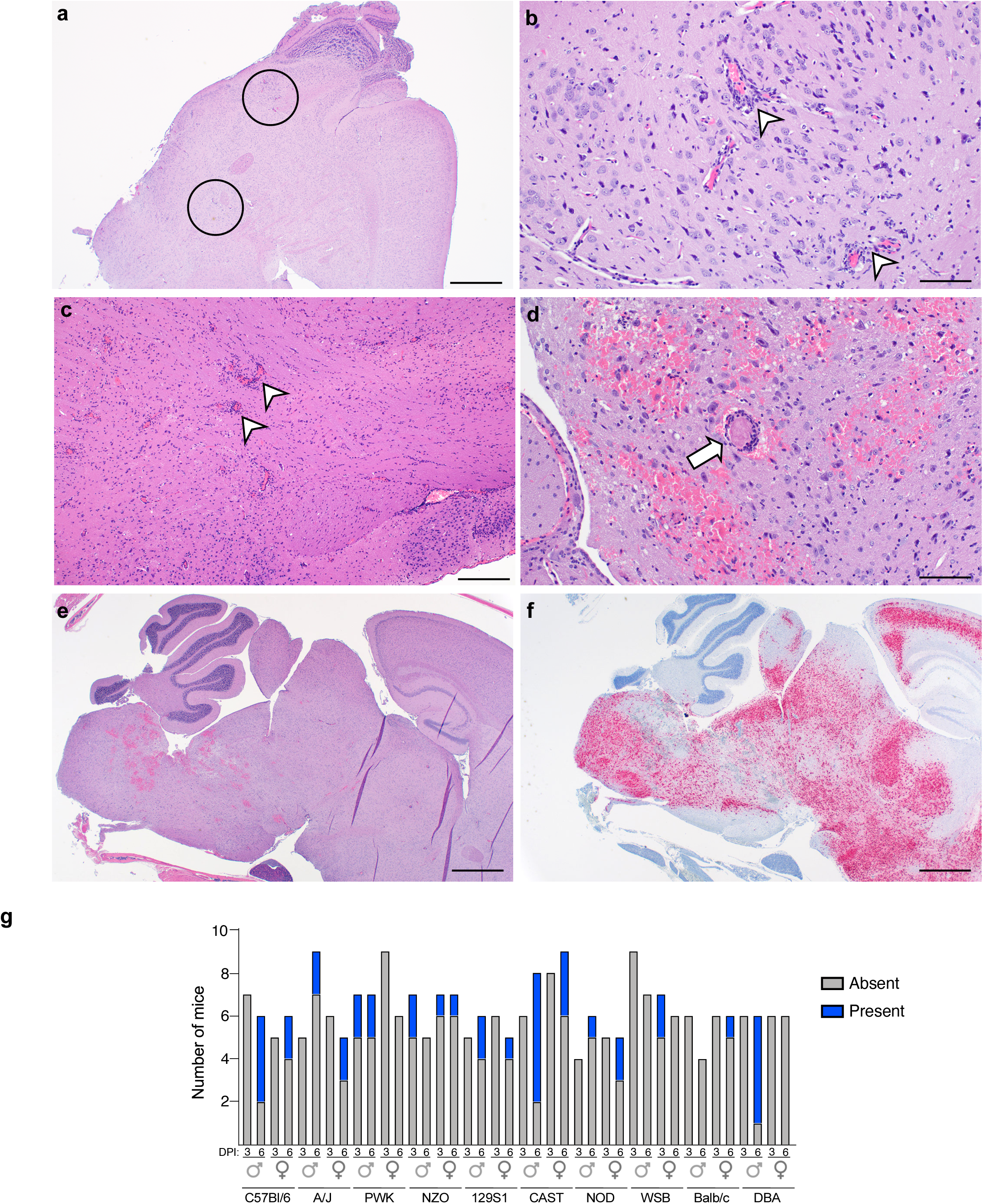
Examples of pathology in the brain of SARS-CoV-2-infected CC x K18-hACE2 F1 progeny. (**a)** Small focal areas of inflammation were commonly observed irrespective of genetic strain (circled). Scale bar = 1000 μm. (**b)** increased magnification of (A) demonstrating perivascular cuffing, gliosis, and evidence of necrotic cells in areas of focal inflammation. Scale bar = 200 μm. (**c)** Example of a pronounced lymphocytic inflammatory response with perivascular cuffing and increased cellularity (arrowheads). Scale bar = 100 μm. **(d)** Microthrombus occluding cerebral capillaries and hemorrhage (arrow) observed in CASTxK18-hACE2. Scale bar = 100 μm. (**e)** Example of extensive hemorrhage in the brain of CASTxK18-hACE2 F1 progeny associated with extensive virus replication as demonstrated in F. Scale bar = 1000 μm. (**f)** serial section relative to **(e)** showing extensive SARS-CoV-2 genome by RNAscope. Scale bar = 1000 μm. **(g)** Incidence of encephalitis for CC Founder F1 males and females at 3 and 6 dpi scored as absent or present.

**Supplemental Figure 4:**
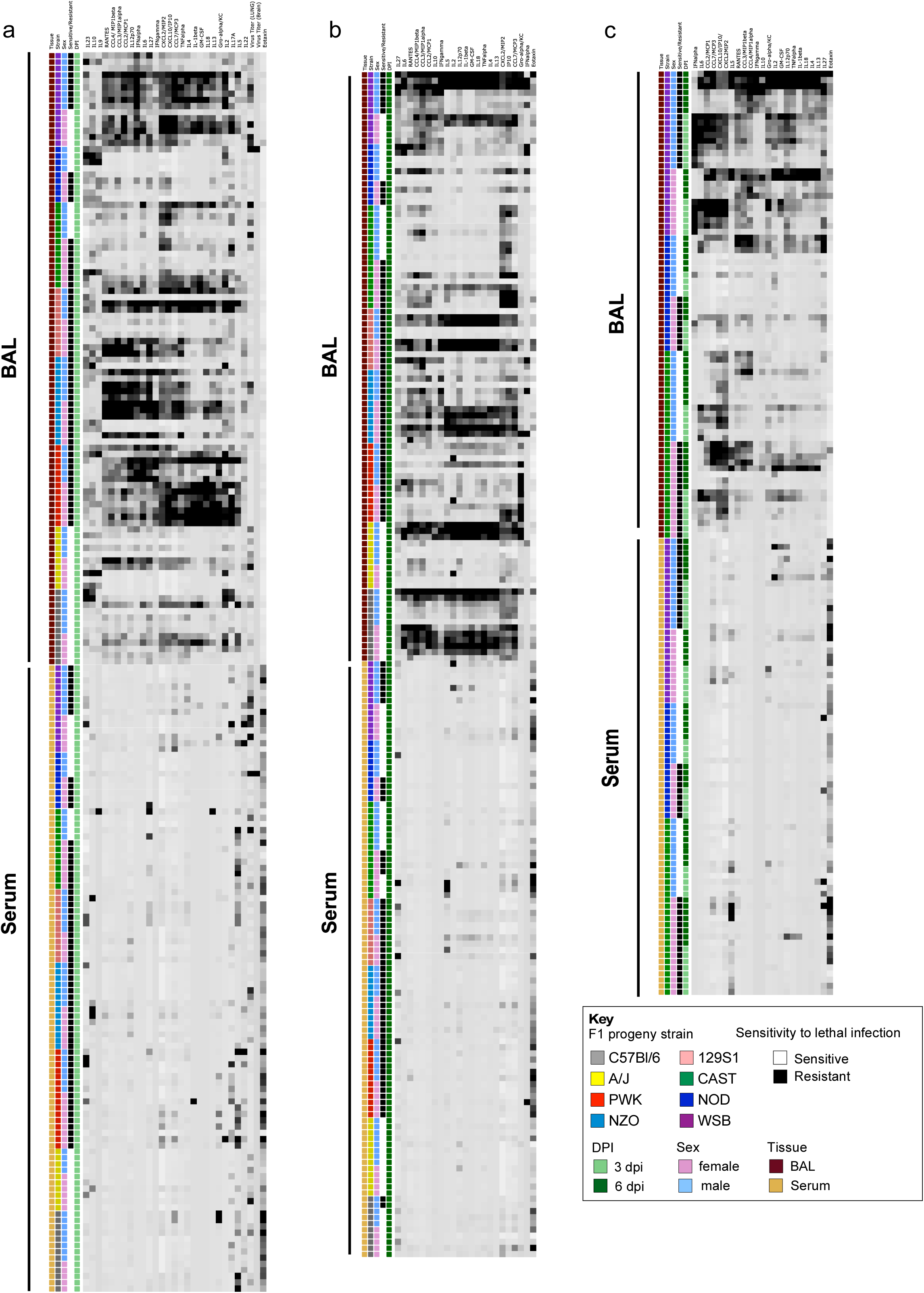
Interferon, cytokine and chemokine expression in the serum compared to the BAL of SARS-CoV-2-infected CC x K18-hACE2 F1 progeny. Heatmap showing relative cytokine levels, as measured by multiplex cytokine assays, for all eight at CCxK18-hACE2 F1s at (a) 3 dpi, and (b) 6 dpi, and (c) for strains with a sex-bias at 3 and 6 dpi.

**Supplemental Figure 5:**
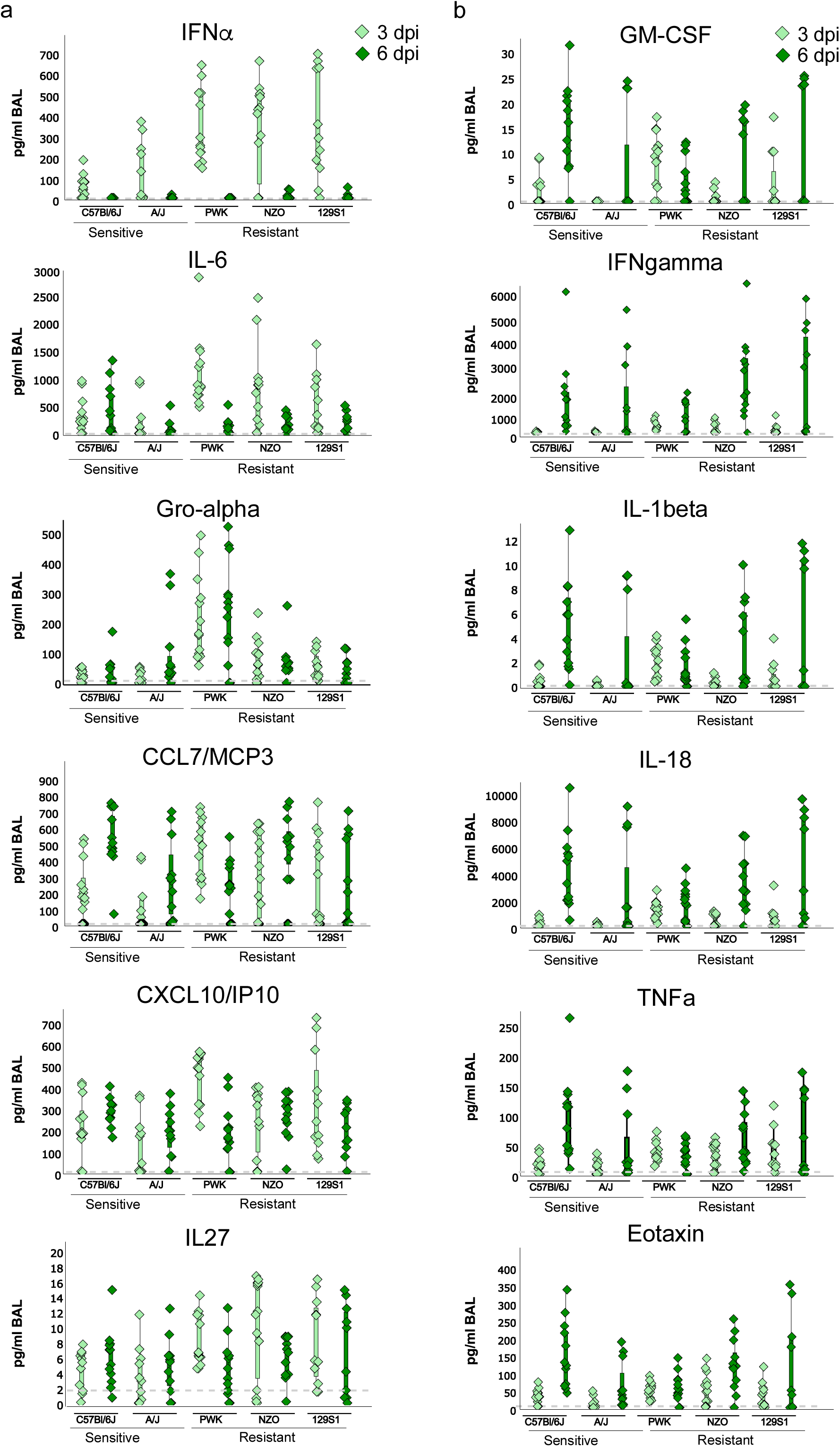
Interferon, cytokine, and chemokine expression in the BAL from SARS-CoV-2- infected sensitive and resistant CC x K18-hACE2 F1 progeny. Cytokines were measured at 3 and 6 dpi by multiplex cytokine assays. Box plots represent data from male and female sensitive (K18-hACE2 and A/J x K18- hACE2) and resistant (PWK, NZO and 129S1J x K18-hACE2) F1s. (a) “Group A” cytokines with higher production in resistant F1 progeny compared to sensitive F1 progeny at 3 dpi. (b) “Group B” cytokines with delayed and sustained production in sensitive compared to resistant F1 progeny. The average cytokine expression in BAL of naïve mice is indicated by a grey dashed line. Box plots (drawn in Qlucore software) show individual data points with the box representing the first and third quartiles.

**Supplemental Figure 6:**
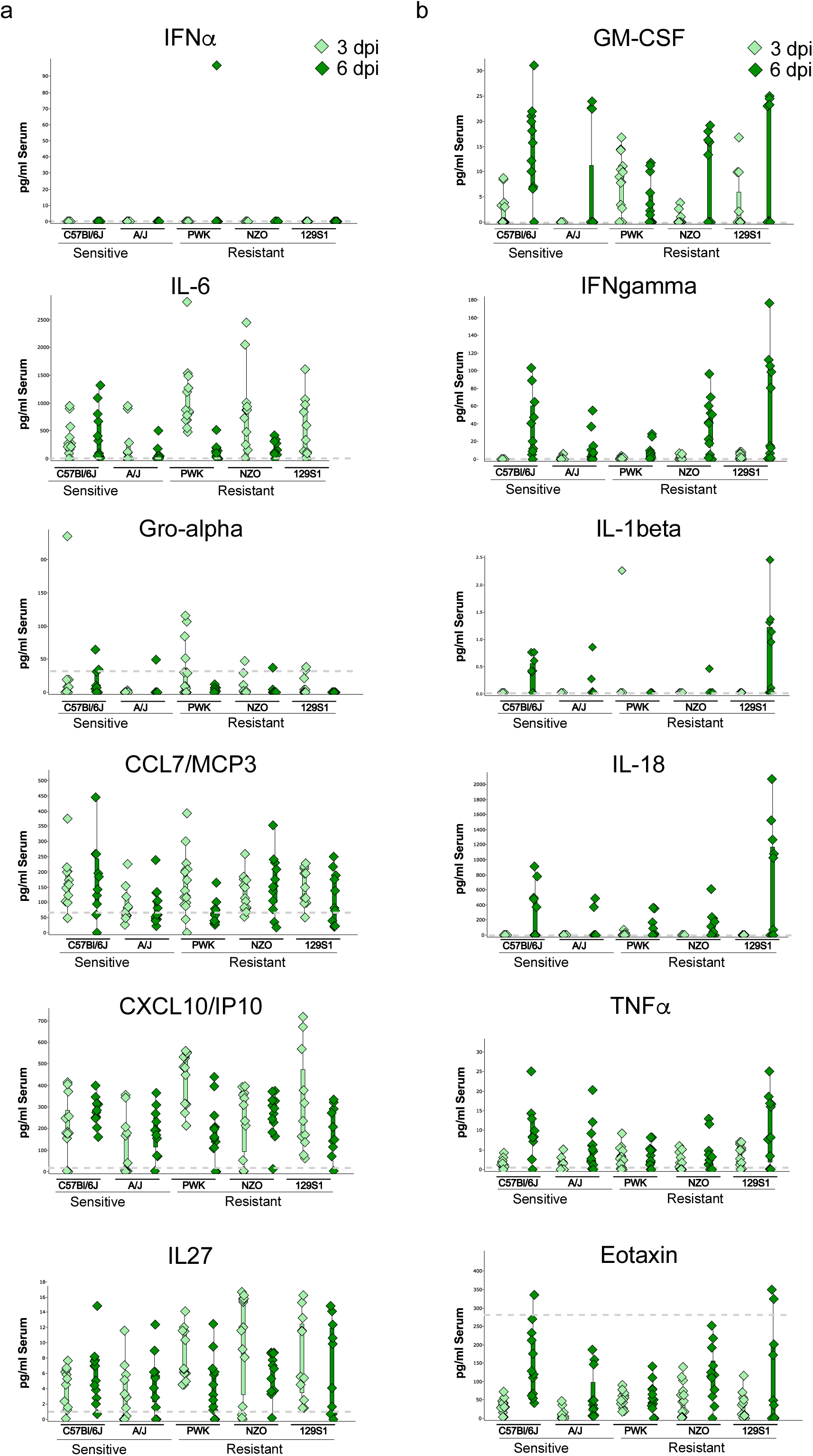
Interferon, cytokine, and chemokine expression in the serum from SARS-CoV-2- infected sensitive and resistant CCxK18-hACE2 F1 progeny. Cytokines were measured at 3 and 6 dpi by multiplex cytokine assays. Box plots represent data from male and female sensitive (K18-hACE2 and A/J x K18- hACE2) and resistant (PWK, NZO and 129S1J x K18-hACE2) F1s. (a) “Group A” cytokines with higher production in resistant F1 progeny compared to sensitive F1 progeny at 3 dpi. The average cytokine expression in serum of naïve mice is indicated by a grey dashed line. Box plots (drawn in Qlucore software) show individual data points with the box representing the first and third quartiles.

**Supplemental Figure 7:**
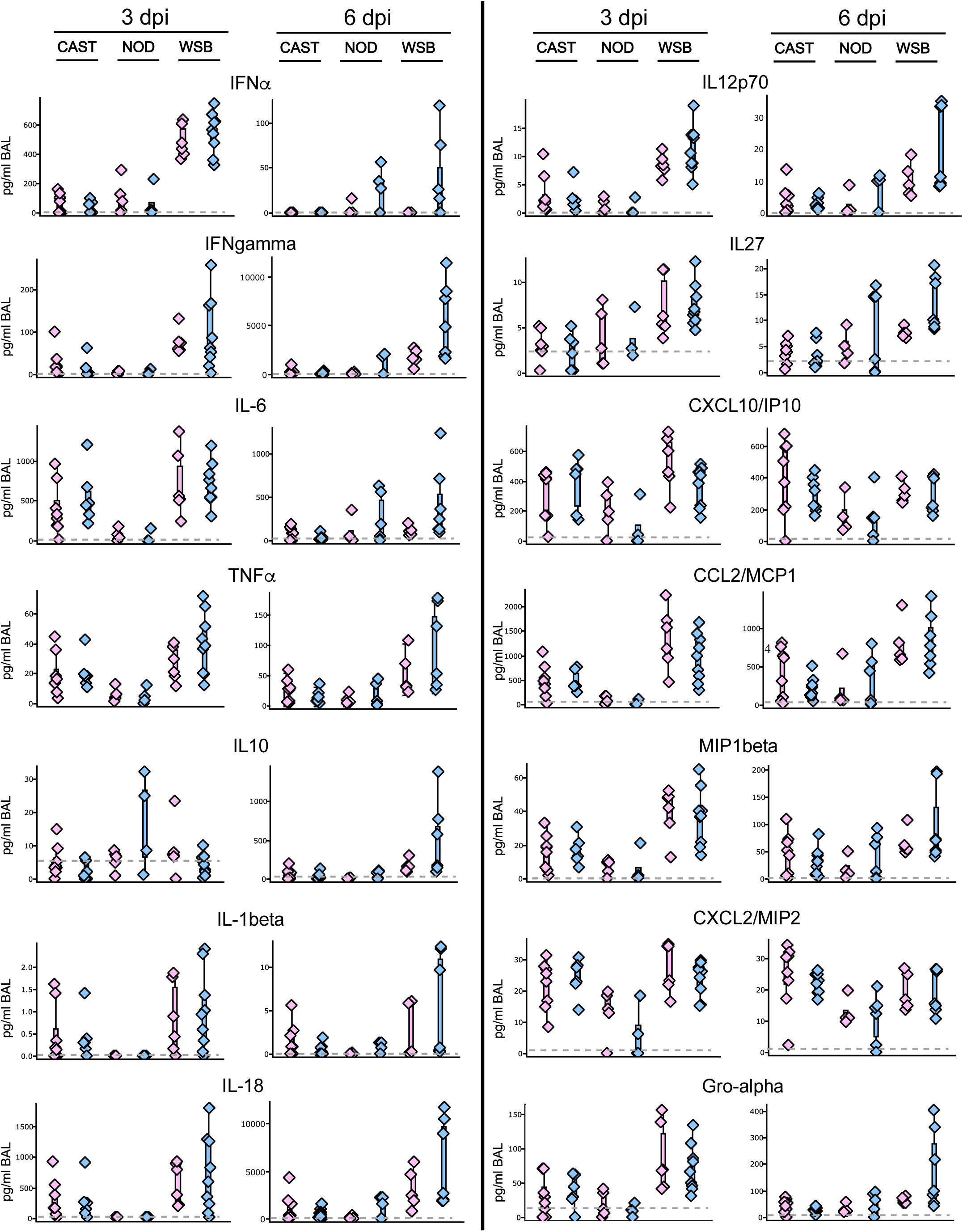
Interferon, cytokine, and chemokine expression in the BAL from SARS-CoV-2- infected CCxK18-hACE2 F1 with a sex bias. Box plots show cytokine levels in male versus female for CAST, NOD, and WSB x K18-hACE2 F1. Cytokines were measured at 3 and 6 dpi by multiplex cytokine assays. The average cytokine expression in BAL of naïve mice is indicated by a grey dashed line. Box plots (drawn in Qlucore software) show individual data points with the box representing the first and third quartiles.

**Table S1.** Summary of percent survival, weight loss, virus burden and BAL cytokine profiles in K18-hACE2 x CC Founder F1 progeny.

| Strain (F1) | Sex | %Survival | Mean time to death (days) | Peak weight loss |  | Average Virus Titers in Lung |  | Cytokine profile BAL |  |
| --- | --- | --- | --- | --- | --- | --- | --- | --- | --- |
|  |  |  |  | % | DPI | 3 dpi Mean (Std Dev) | 6 dpi Mean (Std Dev) | 3 dpi | 6 dpi |
| C57Bl/6 x K18-hACE2 | M | 14 | 6 | 16.7 | 7 | 1.3e6 (1.8e6) | 7.6e4 (1.3e5) | No/Low | High Group A and Group B |
|  | F | 29 | 7 | 12.0 | 7 | 1.5e6 (1.4e6) | 8.2e3 (5.7e3) |  |  |
| A/J x K18-hACE2 | M | 33 | 8 | 15.0 | 7 | 2.8e6 (2.2e6) | 3.5e4 (6.1e4) | No/Low | High Group A and Group B |
|  | F | 66 | UD | 6.1 | 7 | 2.8e6 (2.6e6) | 2.7e4 (2.8e4) |  |  |
| PWK x K18-hACE2 | M | 83 | UD | - | - | 1.6e5 (2.3e5) | BLD | High Group A | Moderate Group A and Group B |
|  | F | 83 | UD | 7.0 | 7 | 9.2e4 (1.1e4) | BLD |  |  |
| NZO x K18-hACE2 | M | 86 | UD | 13.9 | 9 | 2.2e6 (2.8e6) | 3.8e3 (3.7e3) | High Group A | High Group A and Group B |
|  | F | 71 | UD | 21.9 | 9 | 1.4e6 (1.2e6) | 1.3e3 (1.0e3) |  |  |
| 129S1 x K18-ACE2 | M | 86 | UD | 11.8 | 7 | 1.3e6 (7.8e5) | 3.0e3 (2.8e3) | High Group A | High Group A and Group B |
|  | F | 71 | UD | 12.0 | 7 | 8.1e5 (6.4e4) | 2.4e4 (4.0e4) |  |  |
| CAST/EiJ x K18-ACE2 | M | 0 | 6 | 17.2 | 7 | 6.7e6 (7.1e6) | 4.2e5 (3.5e5) | Low Group A | Low Group A |
|  | F | 50 | 15.5 | 35.8 | 8 | 1.6e6 (1.4e6) | 3.4e4 (2.9e4) | Low Group A | Low Group A |
| NOD x K18-ACE2 | M | 33 | 6.5 | 6.0 | 6 | 2.5e6 (4.4e6) | 4.8e3 (5.3e3) | No/Low | Moderate I and II |
|  | F | 83 | UD | - | - | 5.1e5 (7.9e5) | 6.4e3 (8.1e3) | No/Low | Moderate I and II |
| WSB x K18-ACE2 | M | 66 | UD | 32.0 | 11 | 4.1e6 (1.8e6) | 3.1e4 (1.6e4) | Moderate Group A | High Group A and B |
|  | F | 0 | 8 | 25.1 | 8 | 4.4e6 (4.1e6) | 2.5e4 (1.2e4) | Moderate Group A | Moderate Group A and Group B |
| BALB/c x K18-ACE2 | M | 100 | UD | 3.0 | 5 | 6.0e5 (5.5e5) | 1.6e4 (1.3e4) | No/Low | Low Group B |
|  | F | 66 | UD | 8.3 | 6 | 3.7e5 (4.2e5) | 6.0e2 (1.2e3) | High Group A | High Group A and Group B |
| DBA/2 x K18-ACE2 | M | 57 | UD | 16.0 | 10 | 1.5e6 (1.7e6) | 1.7e4 (2.2e4) | No/Low | No/Low |
|  | F | 71 | UD | 3.6 | 6 | 1.7e6 (1.2e6) | 1.2e3 (1.0e3) | No/Low | High Group A and Group B |

undetermined BLD: below limit of detection

**Table S2.**
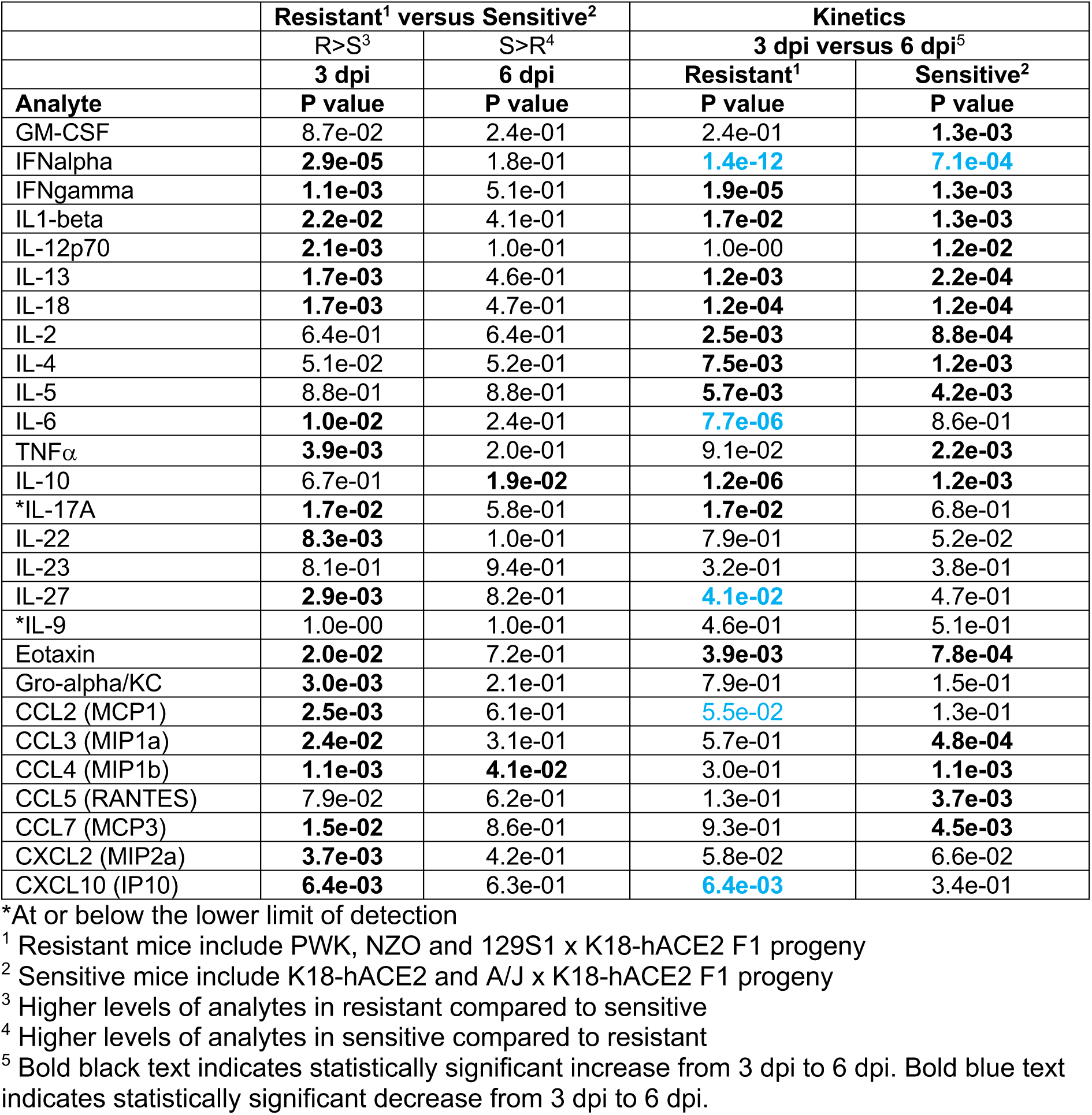
Summary of statistical analyses of cytokine and chemokine expression in BAL fluid collected from resistant and sensitive K18-hACE2 x CC Founder F1 progeny.

| Analyte | Resistant<sup>1</sup> versus Sensitive<sup>2</sup> |  | Kinetics |  |
| --- | --- | --- | --- | --- |
|  | R>S<sup>3</sup> | S>R<sup>4</sup> | 3 dpi versus 6 dpi<sup>5</sup> |  |
|  | 3 dpi | 6 dpi | Resistant<sup>1</sup> | Sensitive<sup>2</sup> |
|  | P value | P value | P value | P value |
| GM-CSF | 8.7e-02 | 2.4e-01 | 2.4e-01 | **1.3e-03** |
| IFNalpha | **2.9e-05** | 1.8e-01 | **1.4e-12** | **7.1e-04** |
| IFNgamma | **1.1e-03** | 5.1e-01 | **1.9e-05** | **1.3e-03** |
| IL1-beta | **2.2e-02** | 4.1e-01 | **1.7e-02** | **1.3e-03** |
| IL-12p70 | **2.1e-03** | 1.0e-01 | 1.0e-00 | **1.2e-02** |
| IL-13 | **1.7e-03** | 4.6e-01 | **1.2e-03** | **2.2e-04** |
| IL-18 | **1.7e-03** | 4.7e-01 | **1.2e-04** | **1.2e-04** |
| IL-2 | 6.4e-01 | 6.4e-01 | **2.5e-03** | **8.8e-04** |
| IL-4 | 5.1e-02 | 5.2e-01 | **7.5e-03** | **1.2e-03** |
| IL-5 | 8.8e-01 | 8.8e-01 | **5.7e-03** | **4.2e-03** |
| IL-6 | **1.0e-02** | 2.4e-01 | **7.7e-06** | 8.6e-01 |
| TNFα | **3.9e-03** | 2.0e-01 | 9.1e-02 | **2.2e-03** |
| IL-10 | 6.7e-01 | **1.9e-02** | **1.2e-06** | **1.2e-03** |
| *IL-17A | **1.7e-02** | 5.8e-01 | **1.7e-02** | 6.8e-01 |
| IL-22 | **8.3e-03** | 1.0e-01 | 7.9e-01 | 5.2e-02 |
| IL-23 | 8.1e-01 | 9.4e-01 | 3.2e-01 | 3.8e-01 |
| IL-27 | **2.9e-03** | 8.2e-01 | **4.1e-02** | 4.7e-01 |
| *IL-9 | 1.0e-00 | 1.0e-01 | 4.6e-01 | 5.1e-01 |
| Eotaxin | **2.0e-02** | 7.2e-01 | **3.9e-03** | **7.8e-04** |
| Gro-alpha/KC | **3.0e-03** | 2.1e-01 | 7.9e-01 | 1.5e-01 |
| CCL2 (MCP1) | **2.5e-03** | 6.1e-01 | **5.5e-02** | 1.3e-01 |
| CCL3 (MIP1a) | **2.4e-02** | 3.1e-01 | 5.7e-01 | **4.8e-04** |
| CCL4 (MIP1b) | **1.1e-03** | **4.1e-02** | 3.0e-01 | **1.1e-03** |
| CCL5 (RANTES) | 7.9e-02 | 6.2e-01 | 1.3e-01 | **3.7e-03** |
| CCL7 (MCP3) | **1.5e-02** | 8.6e-01 | 9.3e-01 | **4.5e-03** |
| CXCL2 (MIP2a) | **3.7e-03** | 4.2e-01 | 5.8e-02 | 6.6e-02 |
| CXCL10 (IP10) | **6.4e-03** | 6.3e-01 | **6.4e-03** | 3.4e-01 |

\*At or below the lower limit of detection
<sup>1</sup> Resistant mice include PWK, NZO and 129S1 x K18-hACE2 F1 progeny
<sup>2</sup> Sensitive mice include K18-hACE2 and A/J x K18-hACE2 F1 progeny
<sup>3</sup> Higher levels of analytes in resistant compared to sensitive
<sup>4</sup> Higher levels of analytes in sensitive compared to resistant
<sup>5</sup> Bold black text indicates statistically significant increase from 3 dpi to 6 dpi. Bold blue text indicates statistically significant decrease from 3 dpi to 6 dpi.

**Table S3.**
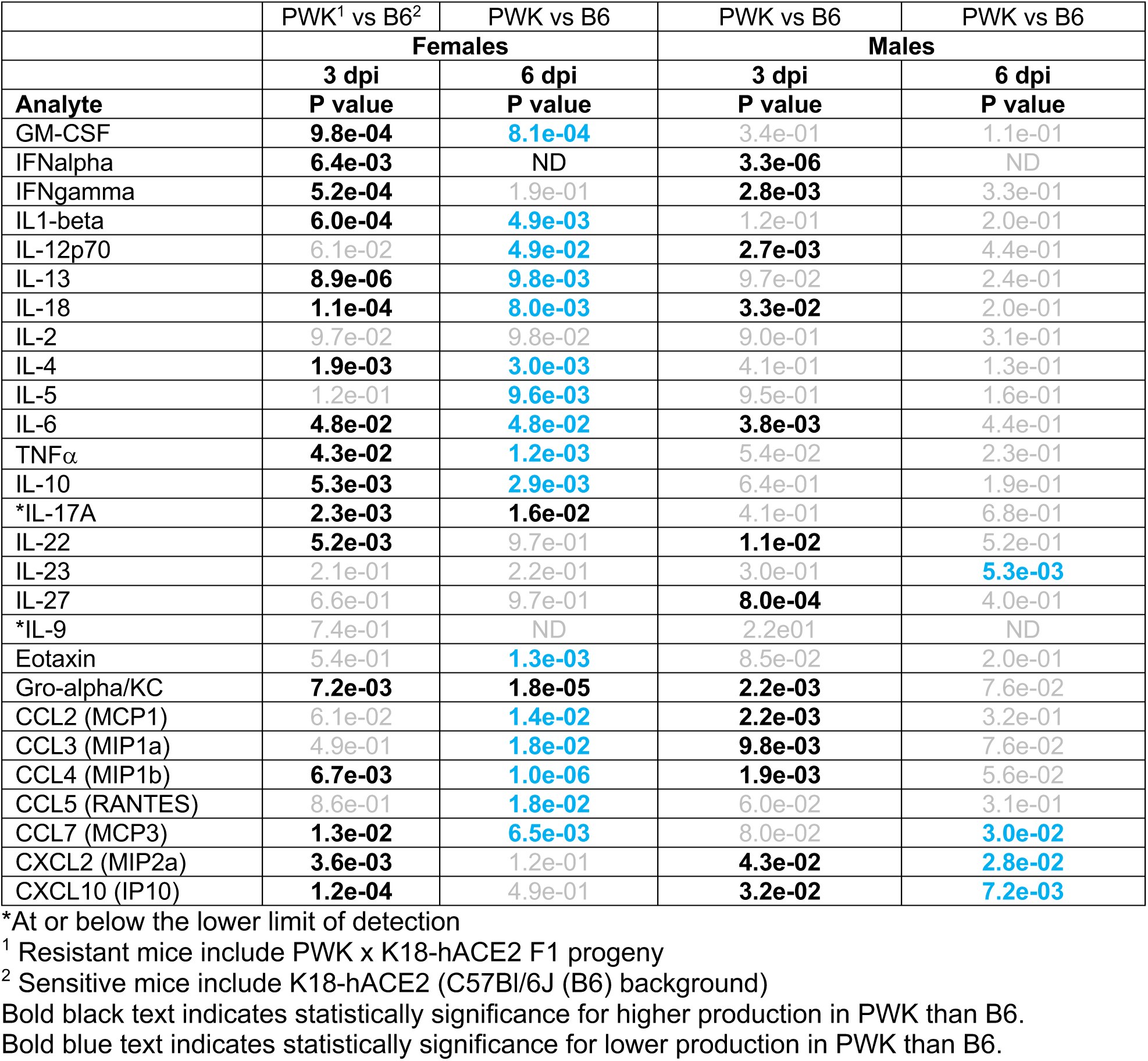
Summary of statistical analyses of cytokine and chemokine expression in BAL fluid comparing male and female resistant PWK x K18-hACE2 and sensitive K18-hACE2 x CC Founder F1 progeny.

## Acknowledgments

This work was funded by the Division of Intramural Research, National Institutes of Health, National Institute of Allergy and Infectious Diseases. Thank you to the animal caretaker staff at campuses of both RML and The Jackson Laboratory for their work in animal husbandry.

## Author contributions

Each author’s contribution to the paper should be listed [we encourage you to follow the CRediT model]. Each CRediT role should have its own line, and there should not be any punctuation in the initials.

Conceptualization: SB, NR, and SMH

Methodology: SJR, NR, SB, and CM

Investigation: SJR, OB, KLM, CS, CC, ML, RMB, AIC, JGS, GLS, RR, SLA, EF, CP, CNB, CP, CB, SM, DPB, JBL, JML, AS, PG, DES, and JS

Visualization: SJR, CNB, JH, DES, CM, DPB, JBL JML, AS, PG, and JS

Funding acquisition: SB and NR

Supervision: SB and NR

Writing – original draft: SB, SJR, CB, AS, NR

Writing – review & editing: SB, SJR, CB, NR

## References

1 Richardson, S. et al. Presenting Characteristics, Comorbidities, and Outcomes Among 5700 Patients Hospitalized With COVID-19 in the New York City Area. JAMA 323, 2052–2059, doi:10.1001/jama.2020.6775 (2020).

2 Long, Q. X. et al. Clinical and immunological assessment of asymptomatic SARS-CoV-2 infections. Nat Med 26, 1200–1204, doi:10.1038/s41591-020-0965-6 (2020).

3 Zhang, Q. et al. Inborn errors of type I IFN immunity in patients with life-threatening COVID-19. Science 370, doi:10.1126/science.abd4570 (2020).

4 Mainous, A. G., Rooks, B. J., Wu, V. & Orlando, F. A. COVID-19 Post-acute Sequelae Among Adults: 12 Month Mortality Risk. Frontiers in Medicine 8, doi:10.3389/fmed.2021.778434 (2021).

5 Docherty, A. B. et al. Features of 20 133 UK patients in hospital with covid-19 using the ISARIC WHO Clinical Characterisation Protocol: prospective observational cohort study. BMJ 369, m1985, doi:10.1136/bmj.m1985 (2020).

6 Pairo-Castineira, E. et al. Genetic mechanisms of critical illness in COVID-19. Nature 591, 92–98, doi:10.1038/s41586-020-03065-y (2021).

7 Wong, L.-Y. R. & Perlman, S. Immune dysregulation and immunopathology induced by SARS-CoV-2 and related coronaviruses — are we our own worst enemy? Nature Reviews Immunology 22, 47–56, doi:10.1038/s41577-021-00656-2 (2022).

8 Schultze, J. L. & Aschenbrenner, A. C. COVID-19 and the human innate immune system. Cell 184, 1671–1692, doi:10.1016/j.cell.2021.02.029 (2021).

9 Park, A. & Iwasaki, A. Type I and Type III Interferons - Induction, Signaling, Evasion, and Application to Combat COVID-19. Cell Host Microbe 27, 870-878, doi:10.1016/j.chom.2020.05.008 (2020).

10 Lucas, C. et al. Longitudinal analyses reveal immunological misfiring in severe COVID-19. Nature 584, 463–469, doi:10.1038/s41586-020-2588-y (2020).

11 Acharya, D., Liu, G. & Gack, M. U. Dysregulation of type I interferon responses in COVID-19. Nature Reviews Immunology 20, 397–398, doi:10.1038/s41577-020-0346-x (2020).

12 Bastard, P. et al. Autoantibodies against type I IFNs in patients with life-threatening COVID-19. Science 370, doi:10.1126/science.abd4585 (2020).

13 Hadjadj, J. et al. Impaired type I interferon activity and inflammatory responses in severe COVID-19 patients. Science 369, 718–724, doi:10.1126/science.abc6027 (2020).

14 Galani, I. E. et al. Untuned antiviral immunity in COVID-19 revealed by temporal type I/III interferon patterns and flu comparison. Nat Immunol 22, 32–40, doi:10.1038/s41590-020-00840-x (2021).

15 Sun, J. et al. Generation of a Broadly Useful Model for COVID-19 Pathogenesis, Vaccination, and Treatment. Cell 182, 734–743.e735, doi:10.1016/j.cell.2020.06.010 (2020).

16 Israelow, B. et al. Mouse model of SARS-CoV-2 reveals inflammatory role of type I interferon signaling. J Exp Med 217, doi:10.1084/jem.20201241 (2020).

17 Gan, E. S. et al. A mouse model of lethal respiratory dysfunction for SARS-CoV-2 infection. Antiviral Res 193, 105138, doi:10.1016/j.antiviral.2021.105138 (2021).

18 Munoz-Fontela, C. et al. Animal models for COVID-19. Nature 586, 509–515, doi:10.1038/s41586-020-2787-6 (2020).

19 Winkler, E. S. et al. SARS-CoV-2 infection of human ACE2-transgenic mice causes severe lung inflammation and impaired function. Nat Immunol 21, 1327–1335, doi:10.1038/s41590-020-0778-2 (2020).

20 Chen, R. E. et al. In vivo monoclonal antibody efficacy against SARS-CoV-2 variant strains. Nature 596, 103–108, doi:10.1038/s41586-021-03720-y (2021).

21 Johnson, B. A. et al. Loss of furin cleavage site attenuates SARS-CoV-2 pathogenesis. Nature 591, 293–299, doi:10.1038/s41586-021-03237-4 (2021).

22 VanBlargan, L. et al. A potently neutralizing anti-SARS-CoV-2 antibody inhibits variants of concern by binding a highly conserved epitope. bioRxiv, doi:10.1101/2021.04.26.441501 (2021).

23 Yinda, C. K. et al. K18-hACE2 mice develop respiratory disease resembling severe COVID-19. PLoS Pathog 17, e1009195, doi:10.1371/journal.ppat.1009195 (2021).

24 Zheng, J. et al. COVID-19 treatments and pathogenesis including anosmia in K18-hACE2 mice. Nature 589, 603-607, doi:10.1038/s41586-020-2943-z (2021).

25 Roberts, A., Pardo-Manuel de Villena, F., Wang, W., McMillan, L. & Threadgill, D. W. The polymorphism architecture of mouse genetic resources elucidated using genome-wide resequencing data: implications for QTL discovery and systems genetics. Mamm Genome 18, 473–481, doi:10.1007/s00335-007-9045-1 (2007).

26 Srivastava, A. et al. Genomes of the Mouse Collaborative Cross. Genetics 206, 537–556, doi:10.1534/genetics.116.198838 (2017).

27 Williamson, E. J. et al. Factors associated with COVID-19-related death using OpenSAFELY. Nature 584, 430–436, doi:10.1038/s41586-020-2521-4 (2020).

28 Ashbrook, D. G., et al. A platform for experimental precision medicine: The extended BXD mouse family. Cell Syst 12, 235-247 e239, doi:10.1016/j.cels.2020.12.002 (2021).

29 Kumari, P. et al. Neuroinvasion and Encephalitis Following Intranasal Inoculation of SARS-CoV-2 in K18-hACE2 Mice. Viruses 13, doi:10.3390/v13010132 (2021).

30 Oladunni, F. S. et al. Lethality of SARS-CoV-2 infection in K18 human angiotensin-converting enzyme 2 transgenic mice. Nat Commun 11, 6122, doi:10.1038/s41467-020-19891-7 (2020).

31 Ackermann, M. et al. Pulmonary Vascular Endothelialitis, Thrombosis, and Angiogenesis in Covid-19. N Engl J Med 383, 120–128, doi:10.1056/NEJMoa2015432 (2020).

32 Gupta, A. et al. Extrapulmonary manifestations of COVID-19. Nat Med 26, 1017–1032, doi:10.1038/s41591-020-0968-3 (2020).

33 Gu, S. X. et al. Thrombocytopathy and endotheliopathy: crucial contributors to COVID-19 thromboinflammation. Nat Rev Cardiol 18, 194–209, doi:10.1038/s41569-020-00469-1 (2021).

34 Bernardes, J. P. et al. Longitudinal Multi-omics Analyses Identify Responses of Megakaryocytes, Erythroid Cells, and Plasmablasts as Hallmarks of Severe COVID-19. Immunity 53, 1296-1314 e1299, doi:10.1016/j.immuni.2020.11.017 (2020).

35 Schulte-Schrepping, J. et al. Severe COVID-19 Is Marked by a Dysregulated Myeloid Cell Compartment. Cell 182, 1419-1440 e1423, doi:10.1016/j.cell.2020.08.001 (2020).

36 Arunachalam, P. S. et al. Systems biological assessment of immunity to mild versus severe COVID-19 infection in humans. Science 369, 1210–1220, doi:10.1126/science.abc6261 (2020).

37 Tostanoski, L. H. et al. Ad26 vaccine protects against SARS-CoV-2 severe clinical disease in hamsters. Nat Med 26, 1694-1700, doi:10.1038/s41591-020-1070-6 (2020).

38 Chakraborty, S. et al. Proinflammatory IgG Fc structures in patients with severe COVID-19. Nat Immunol 22, 67-73, doi:10.1038/s41590-020-00828-7 (2021).

39 Consiglio, C. R. et al. The Immunology of Multisystem Inflammatory Syndrome in Children with COVID-19. Cell 183, 968-981 e967, doi:10.1016/j.cell.2020.09.016 (2020).

40 Mathew, D. et al. Deep immune profiling of COVID-19 patients reveals distinct immunotypes with therapeutic implications. Science 369, doi:10.1126/science.abc8511 (2020).

41 Shen, B. et al. Proteomic and Metabolomic Characterization of COVID-19 Patient Sera. Cell 182, 59–72 e15, doi:10.1016/j.cell.2020.05.032 (2020).

42 Silvin, A. et al. Elevated Calprotectin and Abnormal Myeloid Cell Subsets Discriminate Severe from Mild COVID-19. Cell 182, 1401–1418 e1418, doi:10.1016/j.cell.2020.08.002 (2020).

43 Takahashi, T. et al. Sex differences in immune responses that underlie COVID-19 disease outcomes. Nature 588, 315-320, doi:10.1038/s41586-020-2700-3 (2020).

44 Wilk, A. J. et al. A single-cell atlas of the peripheral immune response in patients with severe COVID-19. Nat Med 26, 1070-1076, doi:10.1038/s41591-020-0944-y (2020).

45 Martin-Sancho, L. et al. Functional landscape of SARS-CoV-2 cellular restriction. Mol Cell 81, 2656–2668.e2658, doi:10.1016/j.molcel.2021.04.008 (2021).

46 Blanco-Melo, D. et al. Imbalanced Host Response to SARS-CoV-2 Drives Development of COVID-19. Cell 181, 1036-1045 e1039, doi:10.1016/j.cell.2020.04.026 (2020).

47 Zanoni, I. Interfering with SARS-CoV-2: are interferons friends or foes in COVID-19? Curr Opin Virol 50, 119–127, doi:10.1016/j.coviro.2021.08.004 (2021).

48 Ziegler, C. G. K. et al. Impaired local intrinsic immunity to SARS-CoV-2 infection in severe COVID-19. Cell 184, 4713-4733 e4722, doi:10.1016/j.cell.2021.07.023 (2021).

49 Sposito, B. et al. The interferon landscape along the respiratory tract impacts the severity of COVID-19. Cell 184, 4953-4968.e4916, doi:10.1016/j.cell.2021.08.016 (2021).

50 Phillippi, J. et al. Using the emerging Collaborative Cross to probe the immune system. Genes & Immunity 15, 38-46, doi:10.1038/gene.2013.59 (2014).

51 Collin, R., Balmer, L., Morahan, G. & Lesage, S. Common Heritable Immunological Variations Revealed in Genetically Diverse Inbred Mouse Strains of the Collaborative Cross. J Immunol 202, 777–786, doi:10.4049/jimmunol.1801247 (2019).

52 Saichi, M. et al. Single-cell RNA sequencing of blood antigen-presenting cells in severe COVID-19 reveals multi-process defects in antiviral immunity. Nature Cell Biology 23, 538–551, doi:10.1038/s41556-021-00681-2 (2021).

53 Mesner, D. et al. SARS-CoV-2 evolution influences GBP and IFITM sensitivity. Proc Natl Acad Sci U S A 120, e2212577120, doi:10.1073/pnas.2212577120 (2023).

54 Delorey, T. M. et al. COVID-19 tissue atlases reveal SARS-CoV-2 pathology and cellular targets. Nature 595, 107-113, doi:10.1038/s41586-021-03570-8 (2021).

55 Hao, Y. et al. Integrated analysis of multimodal single-cell data. Cell 184, 3573–3587.e3529, doi:10.1016/j.cell.2021.04.048 (2021).

56 Hafemeister, C. & Satija, R. Normalization and variance stabilization of single-cell RNA-seq data using regularized negative binomial regression. Genome Biol 20, 296, doi:10.1186/s13059-019-1874-1 (2019).

57 Stuart, T. et al. Comprehensive Integration of Single-Cell Data. Cell 177, 1888–1902.e1821, doi:10.1016/j.cell.2019.05.031 (2019).

